# System-level regulation of hierarchical transitions in a tumour lineage

**DOI:** 10.64898/2026.01.21.700832

**Authors:** Emma Legait, Charlotte Rulquin, Lauranne Bouteille, Raphaël Clément, Cédric Maurange

## Abstract

The fundamental principles defining how cell state transitions are regulated along a tumour lineage to determine its cellular composition remain unclear. Here, we investigate how such transitions are controlled in a reductionist hierarchical brain tumour model. Quantitative analysis of 3D transition maps revealed that the differentiation of cancer stem cells (CSCs) into transit amplifying progenitors (TAPs) depends on the identity of their immediate neighbours. This is embodied in a transition rule that quantitatively predicts the probability to differentiate as a function of the proportion of TAP neighbours. Integrated into 3D simulations of tumour growth, this rule spontaneously recapitulates spatial segregation of CSCs in clusters and their stable proportion. We further show *in vivo* that CSC clustering protects the CSC pool from depletion driven by TAPs. We identify an EGFR-mediated relay mechanism that propagates the CSC-to-TAP transition across CSC clusters. In CSCs, the LRIG1-like EGFR inhibitor Kekkon1 dampens this propagation, ensuring continuous replenishment of the CSC pool. Collectively, these findings show how local fate regulation drives emergent segregation which in turn constrains CSC differentiation dynamics. This provides a conceptual framework to understand and exploit the tumour’s intrinsic differentiation potential by manipulating the determinants of its system-level features.

## Introduction

Hierarchical tumours are driven by a population of cancer stem cells (CSCs) that possess the ability to both self-renew indefinitely or produce more differentiated progenitors, often referred to as transit amplifying progenitors (TAPs), which retain limited proliferative capacity and eventually exit the cell cycle. These hierarchies can be either rigid or plastic, with CSCs maintaining an equilibrium to sustain tumour expansion(Gil Vazquez et al., 2022; Gupta et al., 2011). However, the fundamental mechanisms governing the maintenance and regulation of the CSC population within growing tumours remain poorly understood.

To gain fundamental insights into the principles of CSC self-renewal and differentiation, genetically tractable and reductionist model systems, allowing precise quantifications and formal description, are essential. *Drosophila* neural stem cells (NSCs), also known as neuroblasts, offer a well-established framework to dissect mechanisms underlying stem cell maintenance, asymmetric division, differentiation and tumorigenesis (Doe, 2017; Homem and Knoblich, 2012; Leclercq and Maurange, 2025). Throughout development, they undergo a limited number of asymmetric divisions to self-renew and generate daughter cells rapidly differentiating in neurons and glia. Alteration of the asymmetric division process or perturbation of neuronal differentiation during early development can lead to NSC amplification and sustained proliferation beyond developmental stages, forming tumours that rapidly kill their host (Caussinus and Gonzalez, 2005; Narbonne-Reveau et al., 2016). Because of their early origin, these *Drosophila* tumours represent a powerful model for paediatric cancers (Maurange, 2020). Consistent with the emerging view that intra-tumoral heterogeneity largely mirrors the recapitulation of cellular developmental hierarchies operating in the tissue of origin (Couturier et al., 2020; Karras et al., 2022; Patel and Yanai, 2024; Patel et al., 2014; Vegliante et al., 2021; Vladoiu et al., 2019), these tumours are composed of NSC-like cells exhibiting features of the various stages of their developmental trajectory (Gaultier et al., 2022; Genovese et al., 2019). Some tumour cells co-express the transcription factor Chinmo and the RNA-binding protein Imp (orthologous to human oncofoetal IGF2BPs (Zhu et al., 2023)), reminiscent of early proliferation-prone embryonic and larval stages. Others co-express the transcription factor Eip93F (E93) and the RNA-binding protein Syncrip (Syp) reminiscent of late differentiation-prone larval and pupal stages before adulthood. In both developmental and tumorigenic contexts, Imp promotes *chinmo* translation, whereas Syp inhibits it. Clonal analysis and genetic manipulations have unveiled a rigid cellular hierarchy in tumours with Chinmo being a defining feature of CSC identity in line with its oncogenic properties (Narbonne-Reveau et al., 2016). In contrast, Syp and E93 co-expression defines a TAP identity, these cells rapidly exiting the cell cycle. Interestingly, proportions of CSCs and TAPs rapidly reach a stable equilibrium suggesting robust underlying regulatory mechanisms (Genovese et al., 2019). However, the regulatory principles that govern the probability for a CSC to self-renew or to transit to a more differentiated TAP state are not known. Whether this probability is the same for all cells, thus “hard-coded” in CSCs, or rather depends on external, contextual factors is an open question. How the spatial organization of CSCs may reflect but also affect this outcome is also unknown. Clarifying these principles is essential for grasping how CSC fate decision mechanisms contribute to tumour growth, heterogeneity, and ultimately treatment resistance.

In this study, we leverage 3D segmentation of the different cell states to directly map the CSC-to-TAP transition in tumours. We show that the transition probability is a quantifiable function of the local environment of CSCs, specifically their interactions with neighbouring TAPs. We feed these findings to 3D simulations of tumour growth, and show that such locally-determined fate can spontaneously lead to self-organized CSC clustering and overall stable heterogeneity between cell types in the tumour. This provides a framework to understand how self-organized clustering allows maintaining CSC pools, which we test numerically and experimentally. Finally, we uncover an unexpected pro-differentiation role of EGFR signalling and unveil how this TAP-mediated signal sweeping through CSC clusters is contained by CSCs to maintain intra-tumoral heterogeneity and continuous growth.

Together, our findings reveal that tumours produce their own differentiation cues that regulate locally CSC fate decisions, inducing the emergence of system-level self-organization. This framework provides a mechanistic link between single-cell differentiation behaviour and hierarchical tumour architecture and composition.

## Results

### A cell-intrinsic transition rate fails to recapitulate CSC spatial clustering in 3D simulations

We used a well-established model of developmental tumours initiated during early larval stages by RNAi-mediated knockdown of *prospero (pros)* from a subset of six larval NSCs in the *Drosophila* ventral nerve cord (*pox*^*n*^*-GAL4, UAS-prospero-RNAi*, referred to as *pox*^*n*^*>pros-RNAi* hereafter) (Figure 1A, S1A,B) (Gaultier et al., 2022; Genovese et al., 2019; Narbonne-Reveau et al., 2016). NSCs continuously divide asymmetrically to self-renew and generate an intermediate progenitor that produces two neurons. Upon *pros* inactivation, the intermediate progenitors are unable to differentiate, leading to rapid reversion to a NSC-like state, amplification and tumorigenesis (Caussinus and Gonzalez, 2005; Narbonne-Reveau et al., 2016) (Figure 1A, S1A-C). In a previous work, we had shown that such tumours follow a rigid hierarchy with robust cell state transition probabilities, allowing tumours in adults to be reproducibly composed of about 15-20% of CSCs (Figure 1B-C), the rest of the tumour being composed of TAPs and non-proliferative cells (NPCs) (Genovese et al., 2019). With the exception of a negligible population of neurons, all tumour cells express the NSC marker Miranda (Mira). Within this population, CSCs express Chinmo and Imp, while TAPs and NPCs express Syp (Figure 1C, S1C) and Eip93F (E93), the latter being enriched in NPCs (Genovese et al., 2019). *In vivo* clonal analysis was used to feed a numerical model of lineage growth that allowed extracting transition (CSC to TAP) and cell cycle exit (TAP to NPC) tumour-average rates (Genovese et al., 2019) (Figure 1B). However, the determinants of CSC differentiation underlying the CSC-to-TAP transition rate *k*_*t*_, which ultimately sustains long-term tumour growth, remained unknown. Given that *Drosophila* NSC tumours are not infiltrated by non-tumour cells such as immune cells (Genovese et al., 2019; Voutyraki et al., 2023), our working hypothesis is that *k*_*t*_ has to stem from the tumour lineage itself, being either cell-intrinsic and the same for all CSCs, or locally determined by extrinsic signals mediated by the cellular neighbourhood constituted by other tumour cells (Figure 1D).

**Figure 1:**
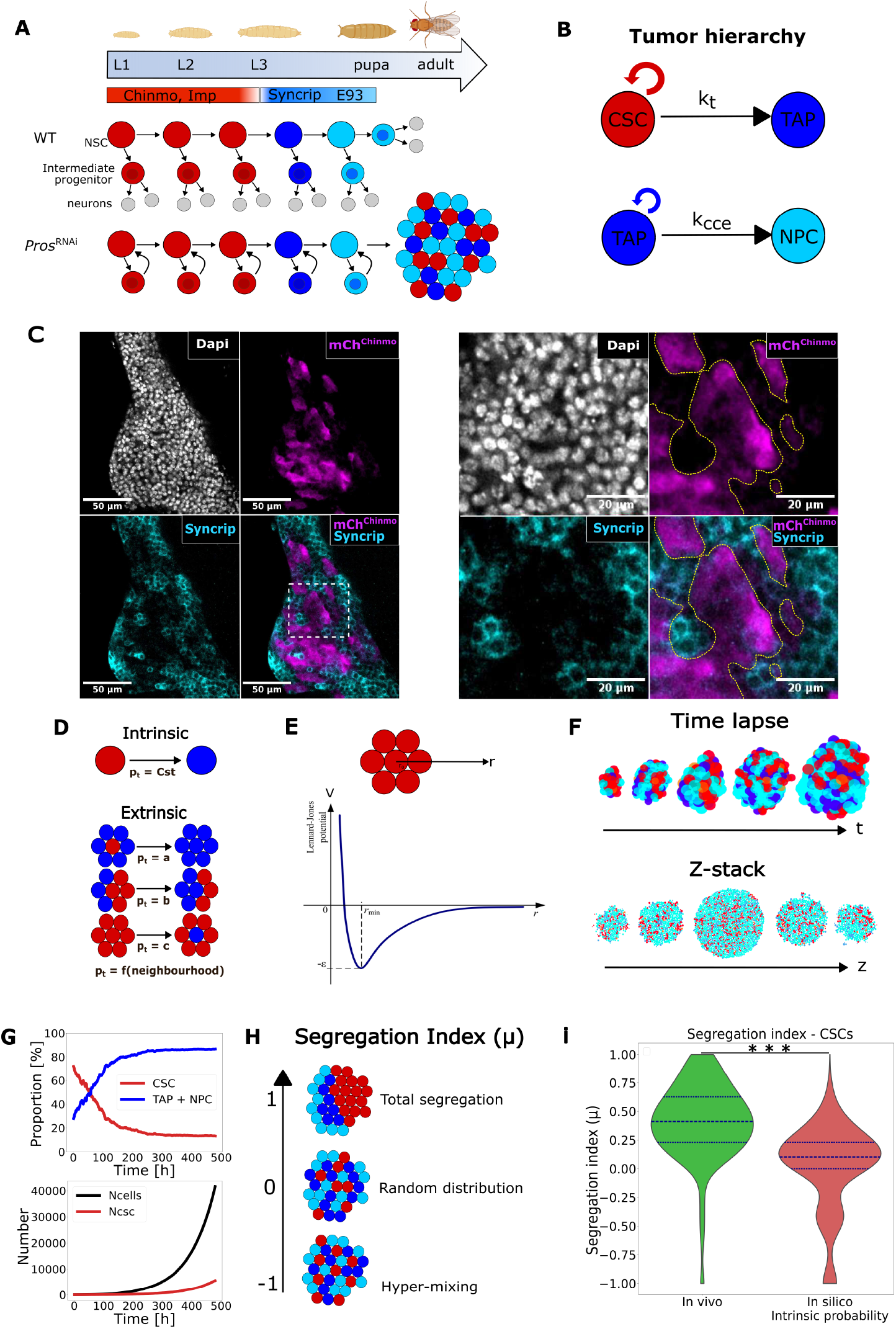
*In vivo* and *in silico* models of hierarchical tumour. **(A)** Asymmetrically-dividing *Drosophila* NSCs transit through three main temporal windows during development defined by Chinmo/Imp (red), Syp (dark blue) and E93 (light blue) respectively. NSCs ultimately differentiate during metamorphosis (pupa). Knockdown of *pros* during the Chinmo/Imp temporal window prevents the differentiation of intermediate progenitors, inducing NSC amplification and tumours that persist in adults. Tumours are composed of NSC-like cells recapitulating the different temporal identities observed during development. **(B)** Tumours exhibit a strict hierarchy with Chinmo^+^ Imp^+^ NSC-like cells behaving as CSCs (red cells), Syp^+^ NSC-like cells behaving as TAPs (dark blue cells) and Syp^+^ E93^+^ cells behaving as NPCs (light blue). *k*_*t*_ is the transition rate of CSCs, and *k*_*cce*_ is the cell cycle exit rate of TAPs. **(C)** A mCherry reporter transgene reflecting chinmo expression (mCh^chinmo^) (see description in Materials and Methods section) is used to label CSCs in tumours. Dapi labels the nucleus of all tumour cells. TAPs + NPCs are labelled with an anti-Syp antibody. Note the clustered organisation of CSCs. The right panels are a magnification of the white dotted square on the left. **(D)** Intrinsic versus extrinsic hypotheses for the regulation of CSC-to-TAP transition. **(E)** Lennard-Jones potential used for the *in silico* model of tumour growth. **(F)** 3D time lapse and Z-stack of a tumour growth simulation. **(G)** Cell proportion over time (top) and number of cells over time (bottom) in a simulation with intrinsically-regulated CSC-to-TAP transition. **(H)** Qualitative description of the segregation index µ. **(I)** CSC segregation index distribution *in vivo*, and *in silico* after simulations with intrinsically-regulated CSC-to-TAP transition (around n = 50 000 cells for both conditions).

To gain insight into how different transition scenarios (intrinsic vs extrinsic) could impact tumour dynamics, we first turned to 3D simulations. We used a particle-based toy model in which each cell is modelled as a hard sphere that physically interacts with its neighbours through a Lennard-Jones potential to mimic the dense cell packing (Figure 1E). Cells can divide (thus pushing their neighbours), undergo CSC-to-TAP transition, or exit the cell cycle according to prescribed rules. The rationale is to use the model as an *in silico* toolbox to make qualitative predictions and make sense of experimental observations. To test the cell-intrinsic scenario, we ran simulations in which all CSCs possess the same transition rate *k*_*t*_ = 0.35 *d*^−1^ and in which TAPs cell cycle exit rate is *k*_*cce*_ = 0.88 *d*^−1^ (Figure 1B,F), consistent with our clonal analysis (see modelling methods for details). Simulations start with 30 CSCs surrounded by 10 TAPs, numbers and spatial organization reminiscent of the early steps of tumour initiation during larval stages (Genovese et al., 2019). CSC and TAPs are set to divide every 1.6 and 1.3 days respectively, as previously determined (Genovese et al., 2019). In these simulations, the overall proportions of CSCs and non-CSCs (TAPs+NPCs, hereafter simply designed as TAPs) reached a plateau (Figure 1G), matching our experimental observations (∼20%/80% respectively). This was expected, as a lineage growth model without space using these probabilities had the same outcome (Genovese et al., 2019). However, CSCs in these 3D simulations appeared randomly distributed throughout the tumour (Figure 1F, Movie S1,S2), contrasting with the clustered organization observed *in vivo* (Figure 1C). To quantify spatial segregation *in silico* and *in vivo*, we computed for each CSC a normalized segregation index µ (Figure 1H). For a given CSC, *µ* = *1* corresponds to perfect segregation, and means that it is surrounded only by other CSCs. By contrast, *µ* = −*1* corresponds to perfect mixing, and means that all surrounding cells are TAPs. Finally, *µ* = *0* corresponds to a balance of TAPs and CSC neighbours that matches the tumour-scale proportions. Thus *µ* = *0* corresponds to a random spatial distribution of cell types. Measuring the distribution of *µ in vivo* using automated 3D segmentation, we found that it is largely superior to 0 (around 0.4 in average), suggesting strong segregation between cell types (Figure 1I). This contrasts with simulations based on the cell-intrinsic transition rate, where segregation was close to 0 (around 0.12 in average), and thus spatial distribution of CSCs close to random. The cell-intrinsic scenario thus fails to account for CSC clustering, which may instead require cell-extrinsic, environment-dependent fate decisions that favour segregation. This view is also supported by the qualitative observation that TAPs are absent from the bulk of CSC clusters (Figure 1C), suggesting that transition does not occur in CSCs surrounded by other CSCs. We thus set out to quantitatively assess *in vivo* whether the transition rate depends on cellular neighbourhood.

### Mapping CSC-to-TAP transitions in tumours reveals local determination of the transition rate

To that end, we first sought to identify and locate cells transitioning from the CSC to the TAP state, thereafter named as transitioning TAPs (tTAPs). During development, NSCs transit from an early Chinmo^+^ Imp^+^ state to a late Syp^+^ state during larval stages (Genovese et al., 2019; Syed et al., 2017). Focusing on the timing of the transition (around early L3), we found that Chinmo is downregulated in NSCs slightly earlier than Imp (Figure 2A). Therefore, during development transitioning NSCs are Chinmo^-^ and Imp^low^. Consistently, we found that a subpopulation of tumour cells did not express Chinmo but retained low levels of Imp. In addition, they exhibited low levels of Syp (Figure 2B). Together, these stainings strongly suggest that this population of cells are tTAPs. Taking advantage of this observation, we used 3D segmentation on confocal stacks coupled to fluorescence quantification to systematically locate CSCs, tTAPs and TAPs, using DAPI to segment nuclei of tumour cells, an mCherry-based Chinmo reporter to distinguish CSCs (Gaultier et al., 2022; Genovese et al., 2019), and Imp immunostaining to identify CSCs and tTAPs. Based on these markers, we defined three cellular categories: (1) Imp^+^ Chinmo^+^ cells as CSCs, (2) Imp^+^ Chinmo^-^ cells as tTAPs, and (3) Imp^-^ Chinmo^-^ cells as TAPs (Figure 2C, S1C-D, Movie S3).

**Figure 2:**
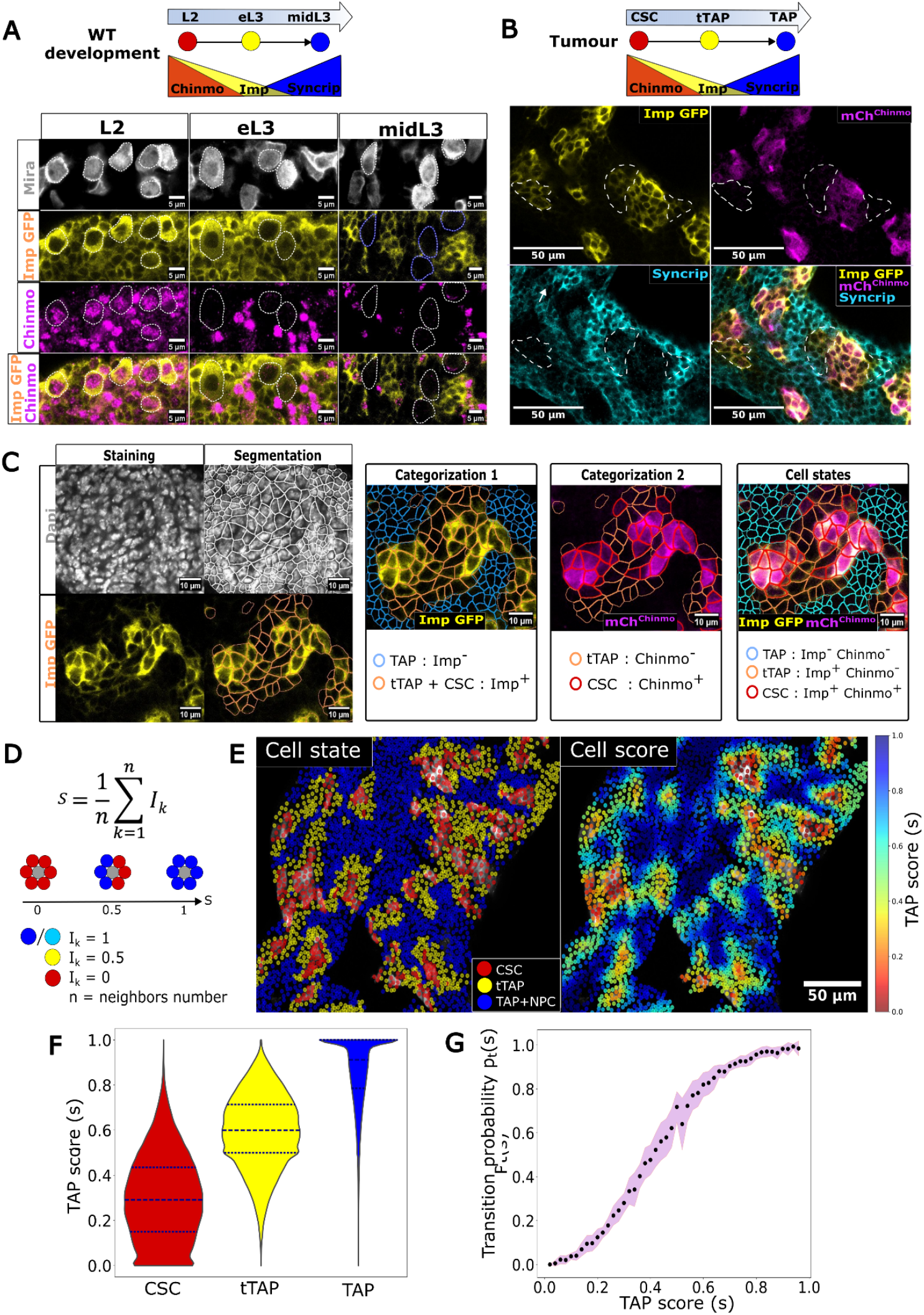
Cell state categorization and transition rule quantification. **(A)** Developmental time course of Chinmo (anti-Chinmo antibody) and Imp expression pattern in NSCs (labelled with Miranda (Mira)) during larval stages revealing a transient Chinmo-Imp+ state in early L3 (eL3). **(B)** A subset of tumour cells (examples delineated by the white dotted lines), exhibits an absence of Chinmo, but low expression of both Imp and Syp suggesting that they are undergoing the CSC-to-TAP transition. **(C)** Segmentation and categorization of cell states: CSC, tTAP, TAP based on mCh^chinmo^, Imp and Dapi stainings. **(D)** Definition of the TAP score *s*. **(E)** 2D sections of the 3D cartography of cell states and TAP scores. **(F)** TAP score distribution in CSCs, tTAPs and TAPs+NPCs. tTAPs have on average more TAP neighbours that CSCs. **(G)** Probability for a CSC to be undergoing transition as a function of the TAP score. This defines a sigmoid-like transition rule (around n = 200 000 cells in total, from 10 different tumours).

With the cell types now mapped, we quantified the local 3D microenvironment of each cell in our tumours using a normalized TAP score *s* typically quantifying the proportion of its neighbours having a TAP identity (Figure 2D). For a given cell, TAP neighbours’ contribution to *s* is 1, and CSC neighbours contribution is 0. We arbitrarily set a 0.5 contribution for tTAPs, halfway between TAPs and CSCs. Combining these 3D maps –cell state and cell score *s* (Figure 2E, Movie S3)– we found that tTAPs have a much higher average proportion of TAP neighbours than other CSCs (Figure 2F). This confirmed that transitions are spatially biased and are more likely to occur in CSCs with more TAP neighbours, typically located at the periphery of CSC clusters. To further characterize the influence of the microenvironment on CSCs fate decisions, we quantified the probability for a CSC to be undergoing transition as a function of the proportion of TAP neighbours, *p*_*t*_ (*s*). This is done by measuring the proportion of tTAPs for different score windows, over tens of thousands of CSCs. We found that *p*_*t*_ (*s*) sharply increases with the proportion of TAPs surrounding them (Figure 2G). CSCs surrounded by other CSCs (i.e. *s* ∼ *0*), have a transition probability close to 0. CSCs located at the periphery of clusters, in contact with both CSCs and TAPs (*s* ∼ *0*.*5*), have an intermediate transition probability. Isolated CSCs (*s* ∼ *1*) are bound to differentiate, with a transition probability close to 1. Interestingly, the sigmoidal transition rule mirrors classical ligand–receptor dose–response curves governed by Hill dynamics (Gesztelyi et al., 2012),hinting towards a possible receptor-mediated mechanism of local fate control. This curve quantitatively relates CSC transition to local neighbourhood composition.

### Stable heterogeneity and segregation emerge from the local transition rule

To investigate the tumour-level implications of this local transition rule, we implemented it directly into our simulation framework. The probability *p*_*t*_(*s*) measured experimentally operates on a timescale *τ* spent by cells in the Chinmo-Imp+ transition state, such that the actual transition rate *k*_*t*_(*s*) to input in simulations is *k*_*t*_(*s*) = *p*_*t*_(*s*)/*τ*. Since *τ* is *a priori* unknown, we conducted a sensitivity analysis for different amplitudes of *k*_*t*_(*s*), corresponding to different values of *τ* (Figure 3A). Simulations implementing the local transition rule showed that the system is able to spontaneously reach stable heterogeneity between cell types (Figure 3A,B). Lower transition rates allow maintaining a stable fraction of CSCs, while higher transition rates lead to CSC population disappearance (Figure 3B). This also ultimately impacts the tumour growth rate (Figure 3C), as the proportion of CSCs is a major determinant of overall tumour growth. In addition, spatial segregation between cell types occurs spontaneously, directly resulting from the local transition rule (Figure 3C). The 15-20% fraction of CSCs observed experimentally is recapitulated for a value of *τ* ≈ 1*d* (Figure 3D, Movie S4). A simple consistency check is to calculate the tumour-average transition rate based on this extrinsic scenario, which should be: 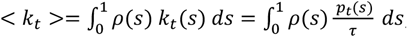, where *ρ*(*s*) is the probability distribution for a CSC to have a score *s*. Using our experimental measurements of *ρ*(*s*), *p*_*t*_(*s*) (Figure 2F,G) and *τ* = 1*d* yields < *k*_*t*_ > ≈ 0.33 *d*^−1^. This is consistent with the value obtained from our clonal analysis (0.35 *d*^−1^). We thus set *τ* = 1*d* in the following simulations.

**Figure 3:**
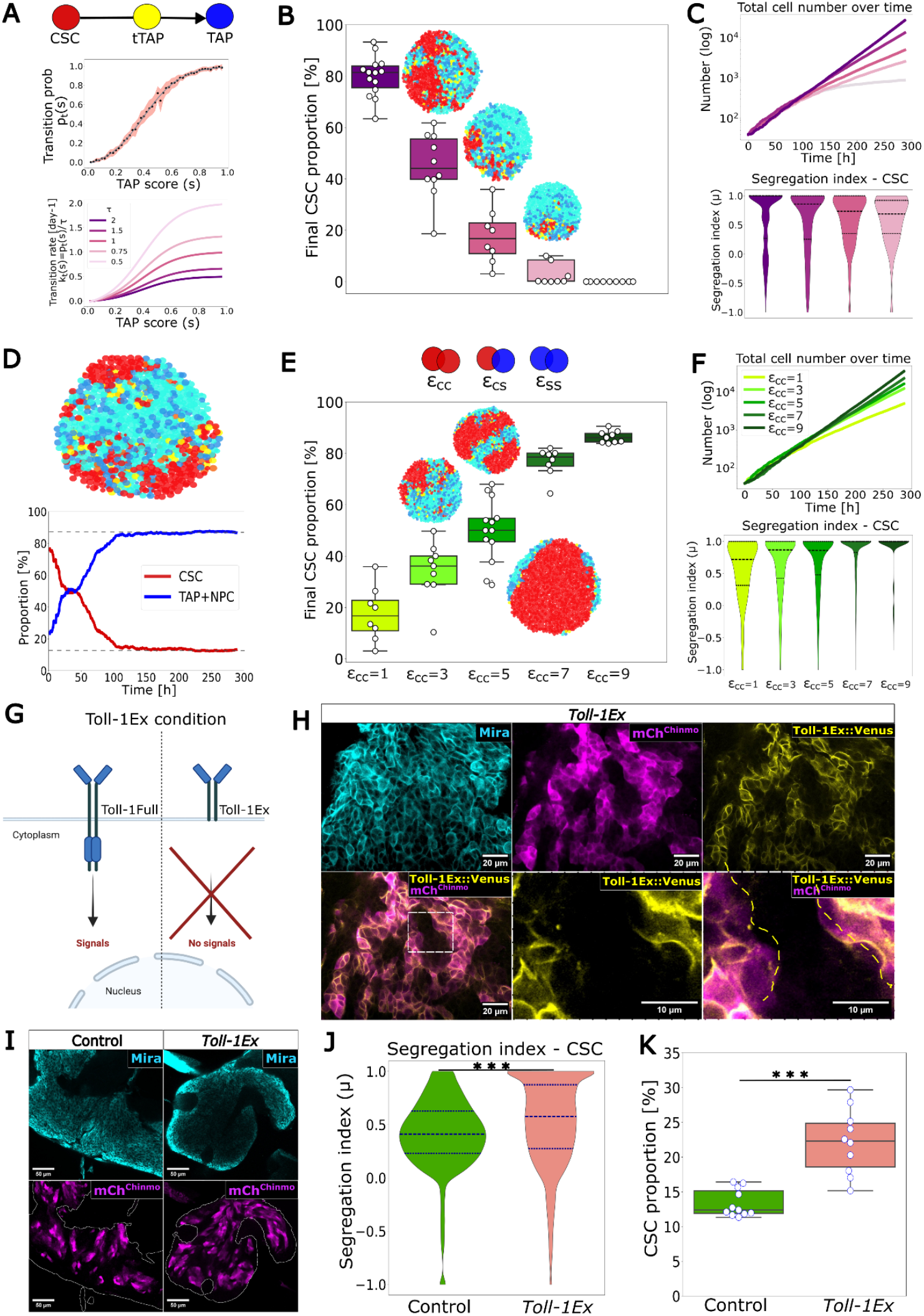
The transition rule drives the emergence of CSC clusters which support CSC maintenance. **(A)** Fit of the experimental transition rule for implementation in the numerical model (top) and resulting transition rates assuming different values of the transition time *τ* (bottom). **(B)** Final proportion of CSCs in simulations (n = 10 per condition) using the transition rates shown in (A). The proportion of CSCs decreases when the average transition rate increases. Colour codes as in (A). **(C)** Cell number over time (top) and segregation index distribution comparison (bottom). Color codes as in (A). **(D)** For a transition rate corresponding to *τ* = *1d*, the system reaches stable heterogeneity and cell type segregation with ∼15% CSCs. **(E)** Final proportion of CSCs in simulations (n=10 per condition) with differential adhesion (*ε*_*cc*_ > *1*). The proportion of CSCs increases when *ε*_*cc*_ increases. **(F)** Cell number over time (top) and segregation index distribution comparison (bottom). Colour codes as in (E). **(G)** Schematic of the *Toll-1Ex* condition. **(H)** Confocal section (40x objective) of the *Toll-1Ex* tumour condition with CSCs labelled with the mCh^chinmo^ reporter transgene and *Toll-1Ex* expression reflected by Venus immune-staining. **(I)** CSC clusters (detected with the mCh^chinmo^ reporter) in control tumours and tumours mis-expressing *Toll-1Ex*. **(J)** Quantification of CSC segregation in Control and *Toll-1Ex* conditions (around n = 60 000 cells for both conditions). **(K)** Quantification of the CSC proportion in Control and *Toll-1Ex* conditions.

Since the transition depends on the proportion of surrounding TAPs, we reasoned that increasing specifically the clustering of CSCs and thus decreasing *ρ*(*s*) should decrease the average transition rate and ultimately increase the overall proportion of CSCs in the tumour. To test this hypothesis numerically, we simply implemented increased adhesion between CSCs to force increased clustering. In our particle-based simulations, cells interact with one another via a Lennard-Jones potential (Figure 1E). By tuning the potential depth *ε*, we can model differential adhesion between cell types. We thus specifically increased the adhesion strength *ε*_*cc*_ between CSCs to mimic stronger homotypic adhesion. As anticipated, we found that stronger adhesion between CSCs led to increased segregation and to a higher stable fraction of CSCs in the tumour (Figure 3E,F, Movie S5), suggesting that selectively changing clustering should be sufficient to impact the CSC fraction and ultimately the overall growth rate (Figure 3F).

Taken together, these simulations show that tumour-scale stable heterogeneity and segregation are emergent properties of the local, cell-extrinsic differentiation rule, and predict that they can be strongly influenced by parameters underlying the transition rate or mechanisms that favour segregation (e.g. differential adhesion). In the following sections, we experimentally test these predictions *in vivo* and investigate the underlying molecular mechanisms.

### CSC clustering promotes their maintenance

To modify *in vivo* the extent of clustering and test the prediction of the model, we engineered a *Drosophila* line in which CSCs specifically overexpress a truncated form of the adhesion protein Toll-1 (lacking the intracellular signalling domain) fused with the Venus GFP variant (Toll-1Ex::Venus) (Figure 3G) (see description in Materials and Methods). Toll-1 has previously been shown to sharpen compartment boundaries by increasing cell-cell adhesion through homotypic interactions in *Drosophila* (Iijima et al., 2020). Since Toll-1 is not expressed in tumours (not detected in single-cell transcriptomic data produced from former studies (Genovese et al., 2019)), this strategy enables us, in principle, to selectively increase adhesive interactions among CSCs without inducing ectopic Toll-1 signalling. Consistently, we observed an expression pattern of the transgene restricted to CSCs, and enrichment of Toll-1Ex-Venus at the interface between adjacent CSCs, compared to the CSC/TAP interface, showing that homotypic interactions stabilize Toll-1Ex at the membrane, as previously described (Iijima et al., 2020) (Figure 3H).

Quantifications demonstrated increased CSC segregation in these tumours (Figure 3I,J), consistent with increased adhesion. Strikingly, this was accompanied by a significant increase in the proportion of CSCs compared to control tumours (Figure 3K). These findings experimentally validate our *in silico* predictions: modifying the cellular microenvironment –here by increasing differential adhesion between cell states— alters the differentiation dynamics by shifting the local transition landscape, leading to an expanded pool of CSCs.

Together, these results demonstrate that CSC clustering is not merely a passive spatial pattern but a constitutive ingredient of stem cell maintenance in tumours where the CSC-to-TAP transition is governed by local cell-cell interactions.

### TAPs induce the CSC-to-TAP transition through activation of EGFR signalling

Given that the CSC-to-TAP transition preferentially occurs at the periphery of CSC clusters and considering the Hill-like transition rule, we hypothesized that a ligand produced by TAPs could signal on CSCs to promote their transition. From single-cell data of *pox*^*n*^>*pros-RNAi* tumours (Genovese et al. 2019), we identified three signalling pathways with receptors and ligands being expressed, known to signal at short distance and commonly involved in cell fate transitions: Notch, Wnt and EGFR (Figure S2A). Interestingly, while knockdown of N, Dl, Fz2 and Wnt5 in all tumour cells did not significantly affect the CSC pool (Figure S2B), we observed a strong increase in the proportion of CSCs upon Spitz knockdown in the tumour (Figure 4A-C). Spitz is a short-range ligand for EGFR, orthologous to mammalian TGFα (Figure 4A). Subsequent knockdown of EGFR or its downstream target ERK also led to a significant increase in CSC proportion, while overexpressing EGFR led to a reduced CSC pool (Figure 4B-C). These results indicate that EGFR activation promotes the transition from CSC to TAP.

**Figure 4:**
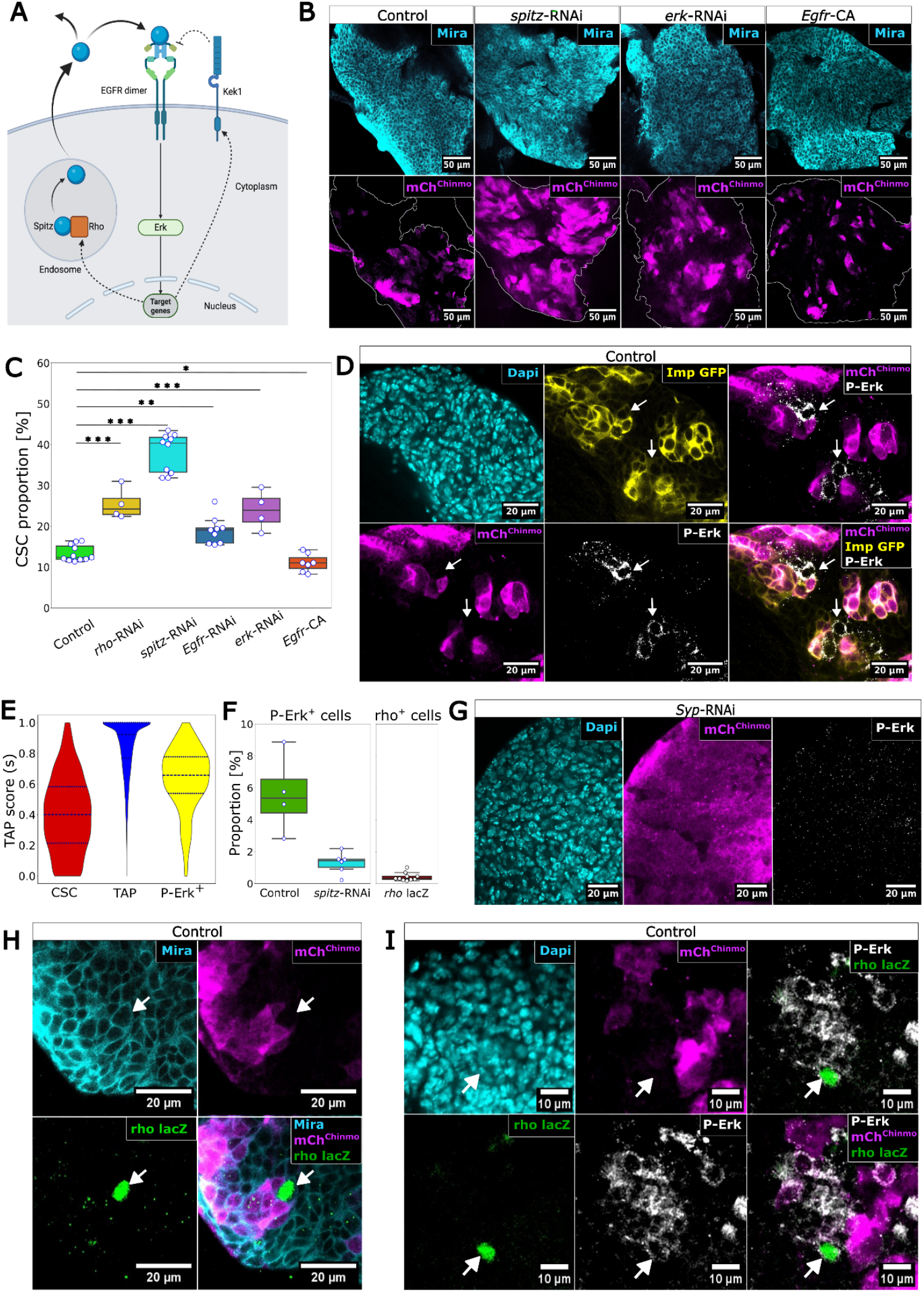
TAP-mediated EGFR signalling promotes CSC differentiation. **(A)** Schematic of the EGFR pathway. **(B)** Mira labels NSC-like tumour cells. mCh^chinmo^ labels CSCs in *control, UAS-spitz-RNAi, UAS-erk-RNAi* and *UAS-Egfr-CA* conditions, confocal section (40x objective). **(C)** Quantification of CSC proportions in the different conditions. **(D)** Confocal section (40x objective) of a Control tumour. Tumour cell nuclei are labelled with Dapi. Arrows indicate examples of anti-P-Erk signal in the cytoplasm of tTAPs identified by lack of mCh^chinmo^ and low expression of Imp-GFP. **(E)** TAP proximity score distribution comparison between CSCs, TAPs and P-Erk+ cells (around n = 20 000 cells). **(F)** Proportion of P-Erk+ cells in control and in *UAS-Spitz-RNAi* conditions (left) and proportion of Rho-lacZ+ cells in control condition (right). **(G)** Confocal section (40x objective) through a *UAS-Syp-RNAi* tumour (only contains CSCs as detected with Chinmo expression) with P-Erk staining. **(H)** Confocal section (40x objective) of control tumour with Rho LacZ staining. **(I)** Confocal section (40x objective) of control tumour with Rho-LacZ and P-Erk stainings.

To investigate the spatial pattern of EGFR pathway activation in the tumour, we monitored phosphorylation of its downstream target ERK with anti-phospho-ERK (P-Erk) immunostaining as a readout. We observed that P-Erk is predominantly detected in the cytoplasm of tTAPs (Chinmo^-^ Imp^+^) (Figure 4D). Statistically, P-Erk^+^ tumour cells exhibit a TAP score lower than TAPs but higher than CSCs (Figure 4E) indicating that they are predominantly located at the periphery of CSC clusters, similar to tTAPs. Consistently, *spitz* knockdown in the tumour decreased the proportion of P-Erk^+^ cells (Figure 4F and S2C). These results support a model in which Spitz produced by cells of the tumour lineage activates EGFR signalling to promote the CSC-to-TAP transition via the MAPK/ERK signalling cascade. To identify which tumour cells produce the EGFR ligand Spitz, we generated tumours composed exclusively of CSCs by silencing Syp (*pox*^*n*^>*pros-RNAi, Syp-RNAi*), and assessed EGFR pathway activity via P-Erk staining. In this context, cytoplasmic P-Erk was undetectable (Figure 4G and S2D). This aligns with a scenario in which committed cells, not CSCs, provide the short-range paracrine signal Spitz that promotes the transition. Supporting this, we detected sparse expression of a *lacZ* reporter gene for *rhomboid* (*rho*), a serine protease that cleaves Spitz to enable its secretion and activation, in a small fraction of P-Erk^+^ tTAPs of control tumours (Figure 4F,H,I). Consistently, quantifying their TAP score *s* indicated that Rho^+^ cells are located at the periphery of clusters (Figure S2E). This suggests that Rho expression may be dynamic and transient in tTAPs, as previously observed in many developmental transitions. Of note, P-Erk was not detected in normal NSCs during larval stages (Figure S2F), indicating that EGFR signalling is not involved in promoting the Imp-to-Syp transition during development. Instead, our results indicate that the regulation of the Imp-to-Syp transition by EGFR signalling is a tumour-specific feature.

Together, these results indicate that differentiation signals originate from newly produced TAPs, consistent with our observation that transition preferentially occurs at the periphery of CSC clusters.

### Kekkon 1 dampens propagation of EGFR-mediated differentiation in CSC clusters

Our simulations reveal that the stable fraction of CSCs is strongly influenced by the transition rate *k*_*t*_, which may be molecularly regulated by factors that modulate EGFR signal intensity within CSCs. Previous work has shown that negative regulators of EGFR signaling (Argos, Kekkon 1; Notch; Sprouty) are deployed in diverse developmental contexts to spatially and temporally fine-tune the outcomes of EGFR-mediated processes (Shilo, 2005). Interestingly, using previously generated single-cell transcriptomic data (Genovese et al., 2019), we identified kekkon 1 (kek1) as one of the most significantly enriched transcripts in CSCs compared to TAPs (Figure 5A). Kek1 encodes for a Leucine-rich repeats (LRRs) and immunoglobulin (Ig) domains (LIG) transmembrane protein, orthologous to LRIG1 in mammals, that is up-regulated in response to EGFR signalling in various *Drosophila* epithelia, and forms a negative feedback loop by binding and inhibiting EGFR (Ghiglione et al., 1999; Ghiglione et al., 2003; Gur et al., 2004). To investigate further *kek1* expression in CSCs, we generated a specific anti-Kek1 antibody, which we confirmed to be functional (Figure S3A). However, immunostaining failed to detect endogenous Kek1 protein in tumours (Figure S3B), pointing towards low or no expression. Nonetheless, forced expression of *kek1* in all tumour cells using the UAS/GAL4 system resulted in a detectable punctate signal throughout the tumour, with notably higher levels in CSCs (Figure S3B,C). This suggests that Kek1 is stabilized in CSCs, consistent with its enriched expression in this population.

**Figure 5:**
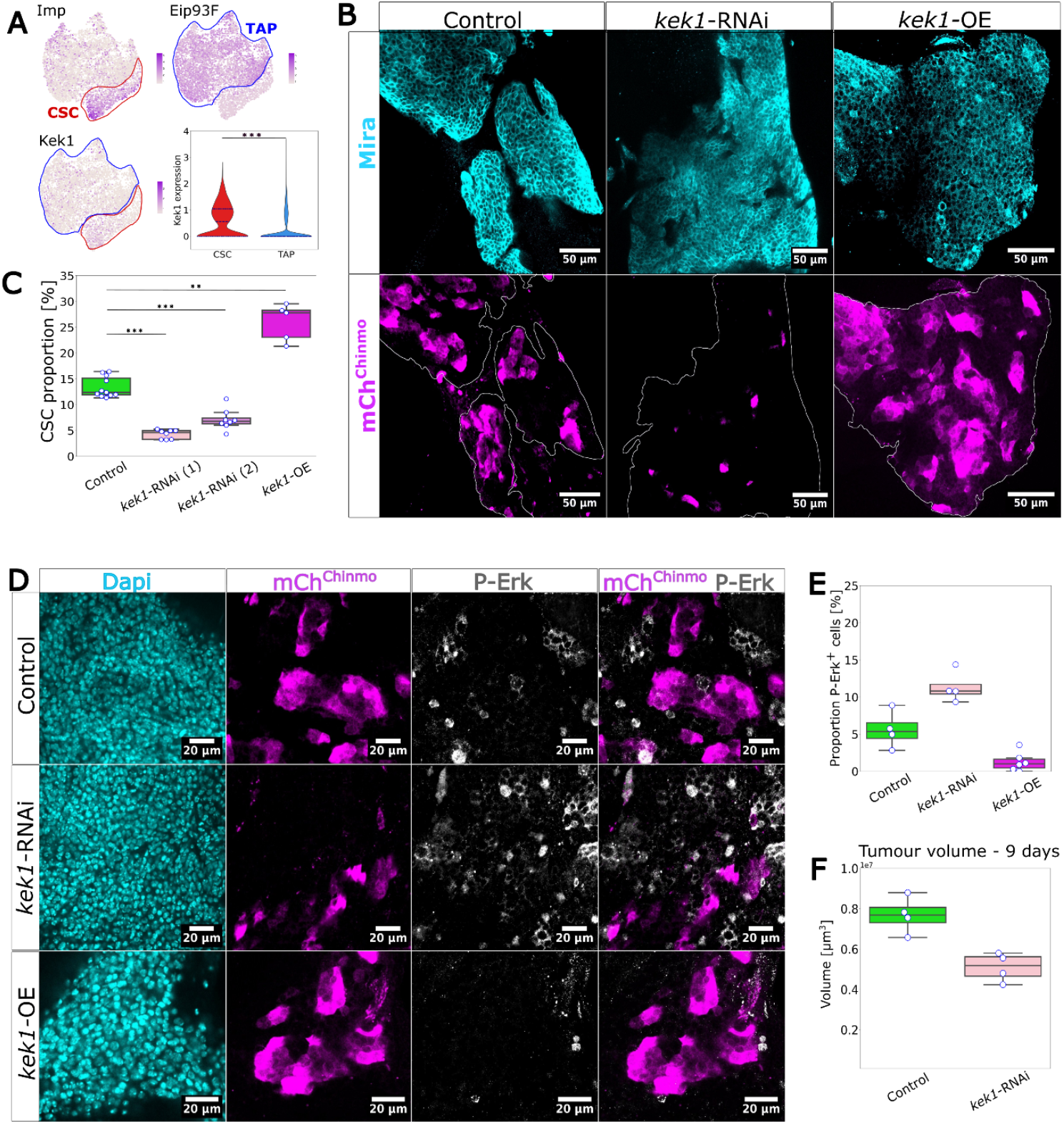
Kek1 expression promotes CSC maintenance by inhibiting the EGFR activation. **(A)** Single cell RNA-seq data with UMAPs of Imp, Eip93F and kek1, and Kek1 expression distribution (bottom right). **(B)** mCh^chinmo^ expression in control, *kek1-* RNAi and *kek1*-OE tumours, confocal section (40x objective). **(C)** Proportion of CSCs in control, *kek1*-RNAi and *kek1*-OE tumours. **(D)** P-Erk expression in control, *kek1*-RNAi and *kek1*-OE tumours, confocal section (40x objective). **(E)** Proportion of P-Erk+ cells in control, *kek1*-RNAi and *kek1*-OE tumours. **(F)** tumour volumes in 9-day-old adults in *control* and *kek1*-RNAi conditions. **(G)** Rho LacZ expression in *kek1*-RNAi tumours, confocal section (40x objective). **(H)** Proportion of Rho+ cells in control and *kek1*-RNAi tumours.

Functionally, *kek1* knockdown led to a marked reduction in the CSC population, indicating that Kek1 is both expressed and functionally active in tumours. Conversely, *kek1* overexpression increased CSC proportion (Figure 5B,C), suggesting that Kek1 promotes CSC maintenance.

To directly test whether Kek1 acts as an EGFR pathway inhibitor, we examined P-Erk levels following *kek1* manipulation. In *kek1* knockdown conditions, P-Erk staining intensity was broadly elevated across tumour cells (Figure 5D,E), indicating increased EGFR signalling. In contrast, tumours overexpressing *kek1* exhibited near-complete loss of P-Erk (Figure 5D,E). Thus, Kek1 inhibits EGFR pathway activation in CSCs. These results support a model in which Kek1 expression renders CSCs less responsive to differentiation activity of the EGFR ligand, hence decreasing the transition rate. Consequently, Kek1 promotes CSC maintenance by favouring CSC self-renewal at the expense of differentiation. Importantly, knockdown of *kek1* led to smaller tumours compared to *control* tumours in 9-day-old adults (Figure 5F) showing that facilitating the CSC-to-TAP transition ultimately triggers depletion of the CSC pool and reduces tumour growth potential.

In conclusion, these findings support a model in which CSC dynamics is governed by paracrine EGFR signalling from newly generated TAPs. The interplay between Spitz, EGFR, Rho, and Kek1 finely tunes the balance between differentiation and maintenance signals that eventually set the transition rate *k*_*t*_ and determines the proportion of CSCs in the tumour (Figure 6A,B).

**Figure 6:**
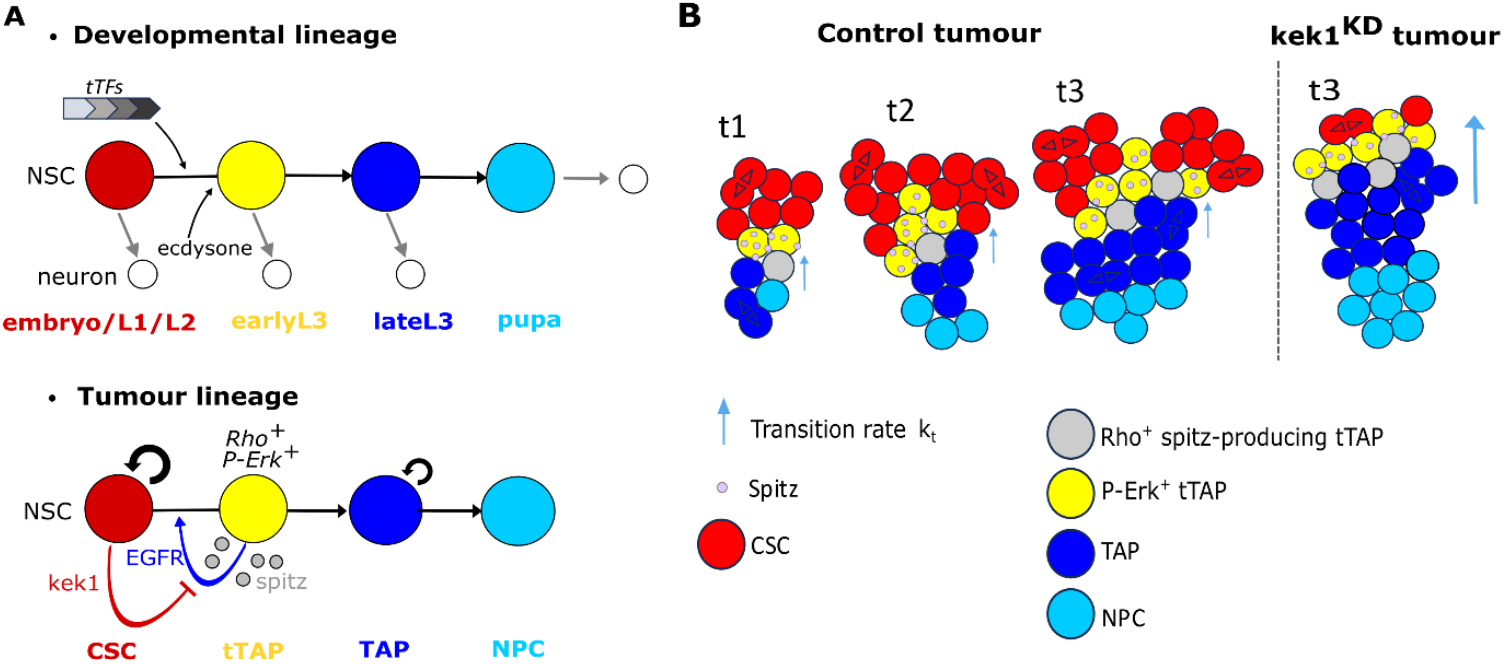
An EGFR-mediated feedback loop regulates CSC population dynamics in the tumour lineage. **(A)** Different regulatory logics for temporal transitions operate in the normal developmental lineage vs tumour lineage. During development, the transition from a Chinmo^+^ Imp^+^ state (in red) to a Syp^+^ state (dark blue) is scheduled in asymmetrically-dividing NSCs by an intrinsic timing mechanism involving the rapid sequential expression of the temporal transcription factors Hunchback→Kruppel→Pou-domain 1→Castor→Grainyhead→Seven-up (Maurange et al., 2008; Ren et al., 2017). In addition, systemic signals like the steroid hormone ecdysone contribute to the transition efficiency (Syed et al., 2017). These mechanisms appear absent in the tumour lineage. Instead, newly generated Syp+ TAPs (tTAPs) produce the EGFR-ligand Spitz that is released to promote the transition to a Syp^+^ TAP state in surrounding Chinmo^+^ Imp^+^ CSCs. CSCs attenuate EGFR signalling by expressing Kek1. **(B)** Schematics of the spatio-temporal dynamics of tumour cell populations upon progression of an EGFR-mediated differentiation front from newly generated TAPs. A balance between differentiation and CSC proliferation maintains population equilibrium and sustains continuous tumour growth. Kek1 knockdown accelerates the transition rate across clusters, resulting in progressive CSC pool depletion and attenuated or arrested tumour growth.

## Discussion

Mammalian tumours form a complex ecosystem in which tumour cells coexist with immune, stromal and vascular infiltrates that generate a rich mixture of chemical and mechanical signals. This complexity makes it difficult to extract overarching rules that govern tumour behaviour at the system level. Using a reductionist approach with a simple hierarchical tumour model in *Drosophila*, we unveil lineage-intrinsic regulatory principles that shape tumour cell heterogeneity. Our findings reveal that the mechanisms underlying the maintenance of a stable CSC proportion are not encoded in individual cells, but are rather a tumour-level outcome of a local differentiation feedback.

By combining computational modelling and genetic experiments on this minimal tumour model, we identify the fundamental mechanisms by which short-range inhibitory interactions between CSCs and their differentiated progeny give rise to self-organized CSC clusters that maintain the CSC fraction within a narrow range. Thus, system-level tumour features emerge from the local interactions underlying hierarchical transitions, and in turn constrain the transition statistics. This mechanism parallels the feedback-mediated regulation of normal stem cell compartments during development and tissue homeostasis, suggesting that tumours exploit conserved principles of tissue self-organization to preserve stemness under dynamic conditions (Dray et al., 2021; Jörg et al., 2019; Lander et al., 2009). Specifically, we demonstrated that the probability for a CSC to transit is a predictable function of its proximity to other TAPs. This transition rule recapitulates the stable heterogeneity observed between cell types, but also the spatial segregation of CSCs, which cannot be explained by a cell-intrinsic differentiation probability. Furthermore, we showed that increasing adhesion between CSCs, both *in silico* and *in vivo* via Toll-1 extracellular domain overexpression, reinforces clustering and promotes CSC amplification. This further confirmed that the local context plays a decisive role in fate determination and that clustering affects differentiation dynamics. Thus, the spatial organization of tumour cells both arises from and constrains differentiation dynamics, thereby regulating the CSC population and the overall tumour growth rate. The sigmoid transition rule is likely to reflect ligand binding statistics governed by Hill equation. This is supported by our results showing the important role of the Spitz-EGFR pathway in the transition process. Our data support a model in which an EGFR-mediated differentiation front progresses throughout CSC clusters to promote transition. Our observations suggest that recently generated TAPs transiently express Rho to release Spitz, propagating EGFR signalling to nearby CSCs leading to sustained differentiation at the front (Figure 6B), a relay model well-described during development (Jörg et al., 2019; Ogura et al., 2018; Yasugi et al., 2010). We found Kek1, a well-characterized EGFR inhibitor, to be expressed at low levels in CSCs. Its activity counteracts EGFR-mediated differentiation, slowing down the progression of differentiation fronts in CSC clusters. Thus, by reducing the transition rate, Kek1 ensures that CSC pools are continuously replenished, tuning the equilibrium between self-renewal and differentiation towards increased self-renewal, a phenomenon predicted by our computational model (Figure 3B).

It is unclear in our tumour context whether Kek1 is rapidly activated in tTAPs in response to EGFR signalling like in other tissues during development (Ghiglione et al., 1999), or constitutively expressed in all CSCs. Interestingly, Kek1 can also strongly bind to itself (MacLaren et al., 2004). Such homotypic interactions may also induce adhesion between CSCs thereby promoting clustering and altering differentiation dynamics by restricting contacts with Spitz-producing TAPs, ultimately sustaining CSC pools. Of note, P-Erk in transitioning cells was mainly cytoplasmic. Interestingly, it has recently been shown in *Drosophila* cell culture, that EGFR/ERK signalling can directly phosphorylate Imp to disassemble the cytoplasmic condensates that they form with their target mRNAs (Graeve et al., 2024). This could lead to inhibition of Imp activity and subsequent chinmo silencing, providing a potential post-translational mechanism by which transient EGFR/ERK signalling triggers the CSC-to-TAP transition. It remains unclear whether EGFR signalling is the only differentiation cue in our *Drosophila* tumour model. The perdurance of some P-Erk levels upon EGFR knockdown associated with an incomplete blockage of the CSC-to-TAP transition suggests either partial inactivation or that other signals may act redundantly.

Our study also sheds light on how developmental programs are co-opted during tumour initiation. In normal development, the Imp-to-Syp transition is precisely scheduled by the coordinated action of neural stem cell-intrinsic timing mechanisms and systemic cues such as hormones (Maurange et al., 2008; Narbonne-Reveau et al., 2016; Syed et al., 2017). Unlike in tumours, we show here that EGFR signalling is not activated in NSCs in the developmental context (Figure 6A). Thus, distinct cues govern the Imp-to-Syp transition in development and in tumorigenesis, with EGFR activation representing a novel trait acquired by the tumour lineage. How such signalling modules arise in neoplasia, rewire developmental trajectories and sustain aberrant proliferation remains a central question in understanding paediatric tumour initiation.

The simplicity of reductionist models is both a strength and a limitation. In our *Drosophila* tumour model, we do not account for potential contributions from immune infiltration—which, although typically low in paediatric cancers, may nonetheless influence TAP-to-CSC feedback dynamics. Additional microenvironmental cues, such as mechanical interactions or steric constraints, may also contribute to shaping tumour organization at the system level. Moreover, long-term *ex vivo* imaging remains technically challenging, limiting our ability to resolve transient pathway activation and differentiation dynamics. Such data could inform numerical models that incorporate more environmental or mechanical cues as well as dynamical aspects of the differentiation signal, notably *rho* expression and EGFR pathway activation. Importantly, these features—absent from our toy model—are likely to affect spatial segregation between cell states, which our model captures qualitatively but not fully quantitatively, as it predicts stronger segregation than observed *in vivo* (Figure 3C). They may also help uncover the determinants of the sigmoid transition rule itself. This outlines a roadmap for future, more quantitative models, both in this system and in progressively more complex experimental contexts. We anticipate that analogous reductionist approaches using human tumoroid-immune co-culture models will help bridge this gap in the future.

Finally, our work challenges the prevailing paradigm in the field that EGFR is uniformly associated with CSC proliferation, survival, and maintenance. However, pro-differentiation roles of EGFR signalling are well-established during development or after seizure in the mammalian brain (Aguirre et al., 2010; Pastor-Alonso et al., 2024). This duality underscores the context-dependent nature of EGFR signalling, where outcomes are shaped by cellular context, signal intensity, and cross-regulation with other pathways, all of which are subject in tumours to ongoing evolution driven by accumulating mutations and therapeutic pressure. Our observations prompt a deeper investigation into the structural and regulatory roles of EGFR signalling in tumour architecture and cell state dynamics at the different stages of tumour progression. They encourage us to explore how the differentiation potential of EGFR could be harnessed therapeutically to drive CSC differentiation.

Here, we revealed a feedback loop between CSC differentiation and tumour spatial organization that determines the proportion of CSCs and the tumour growth rate. Beyond targeting oncogenic drivers, our framework—combining spatial quantification, simulation, and genetic perturbation—suggests that therapeutic value may lie in modulating the tumour’s system-level organization by actively steering the feedback mechanisms that govern the balance between CSC self-renewal and differentiation.

## Supplementary figures

**Supplementary Figure S1:**
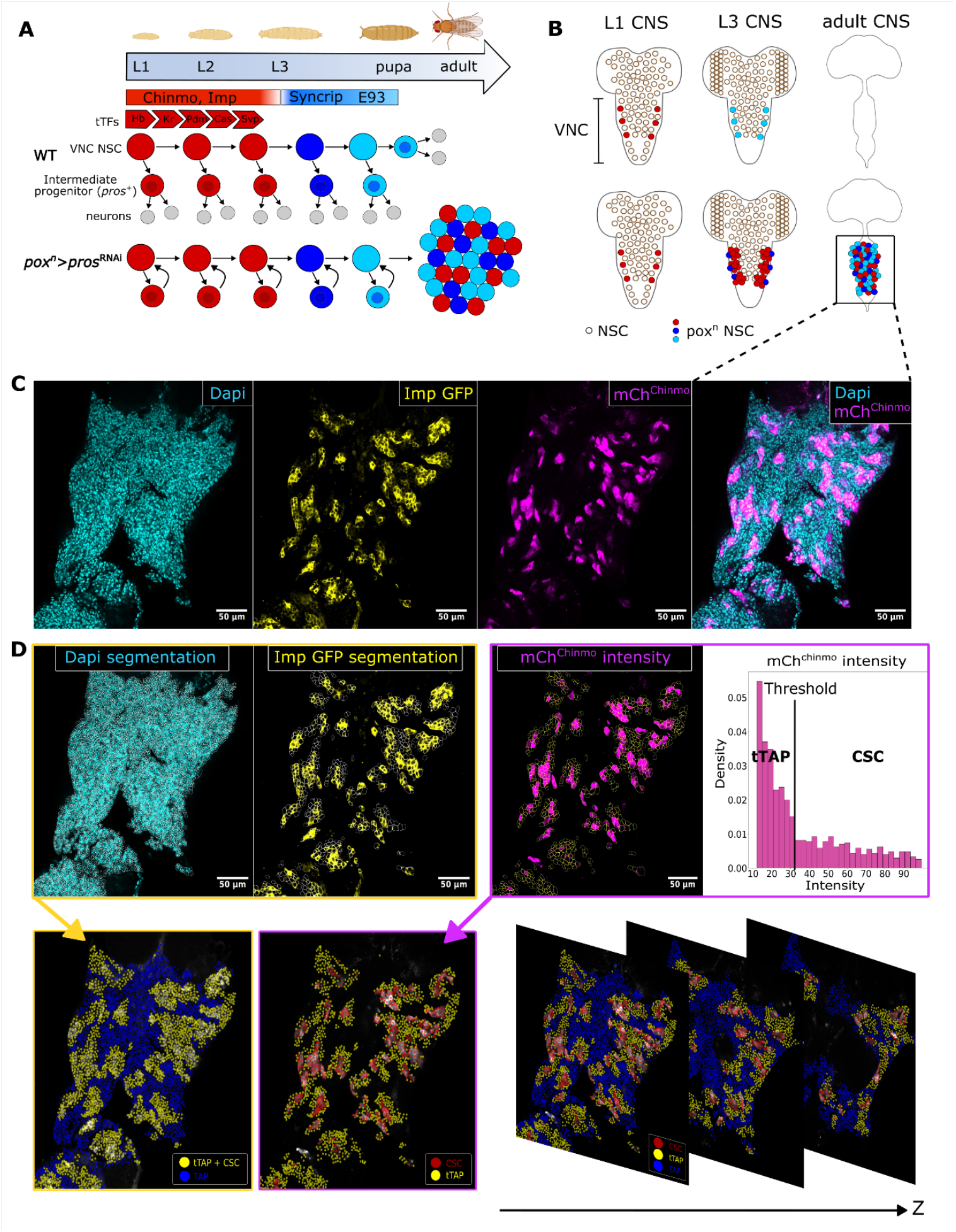
Segmentation pipeline. **(A)** Asymmetrically-dividing *Drosophila* NSCs transit through three main temporal windows during development defined by Chinmo/Imp (red), Syp (dark blue) and E93 (light blue) respectively. NSCs ultimately differentiate during metamorphosis (pupa). During development, the transition from a Chinmo/Imp+ state (in red) to a Syp+ state (dark blue) is scheduled in asymmetrically-dividing NSCs by an intrinsic timing mechanism involving the rapid sequential expression of the temporal transcription factors Hunchback→Kruppel→Pou-domain 1→Castor→Grainyhead→Seven-up (Maurange et al., 2008; Ren et al., 2017). Knockdown of *pros* during the Chinmo/Imp temporal window prevents the differentiation of intermediate progenitors, inducing NSC amplification and tumours that persist in adults. Tumours are composed of NSC-like cells recapitulating the Chinmo/Imp and Syp/E93 temporal identities observed during development. (B) The *pox*^*n*^>*pros*^RNAi^ system allows the generation of tumours from six NSCs of origin located in the VNC. These tumours persist in adults sustained by the Chinmo/Imp+ NSC subpopulation. (C) Tumours persisting in adults can be immunostained and confocal stacks with a 40x objective allow 3D reconstitution with cellular resolution. Dapi shows all the tumour cells and mCh^Chinmo^ show the CSCs. (D) Segmentations of Dapi and Imp stainings allow separating TAPs from the other cell types (tTAPs and CSCs). mCh^Chinmo^ intensity thresholding allows to separate tTAPs from CSCs. This pipeline allows the complete categorization of all tumour cells in 3D.

**Supplementary Figure S2:**
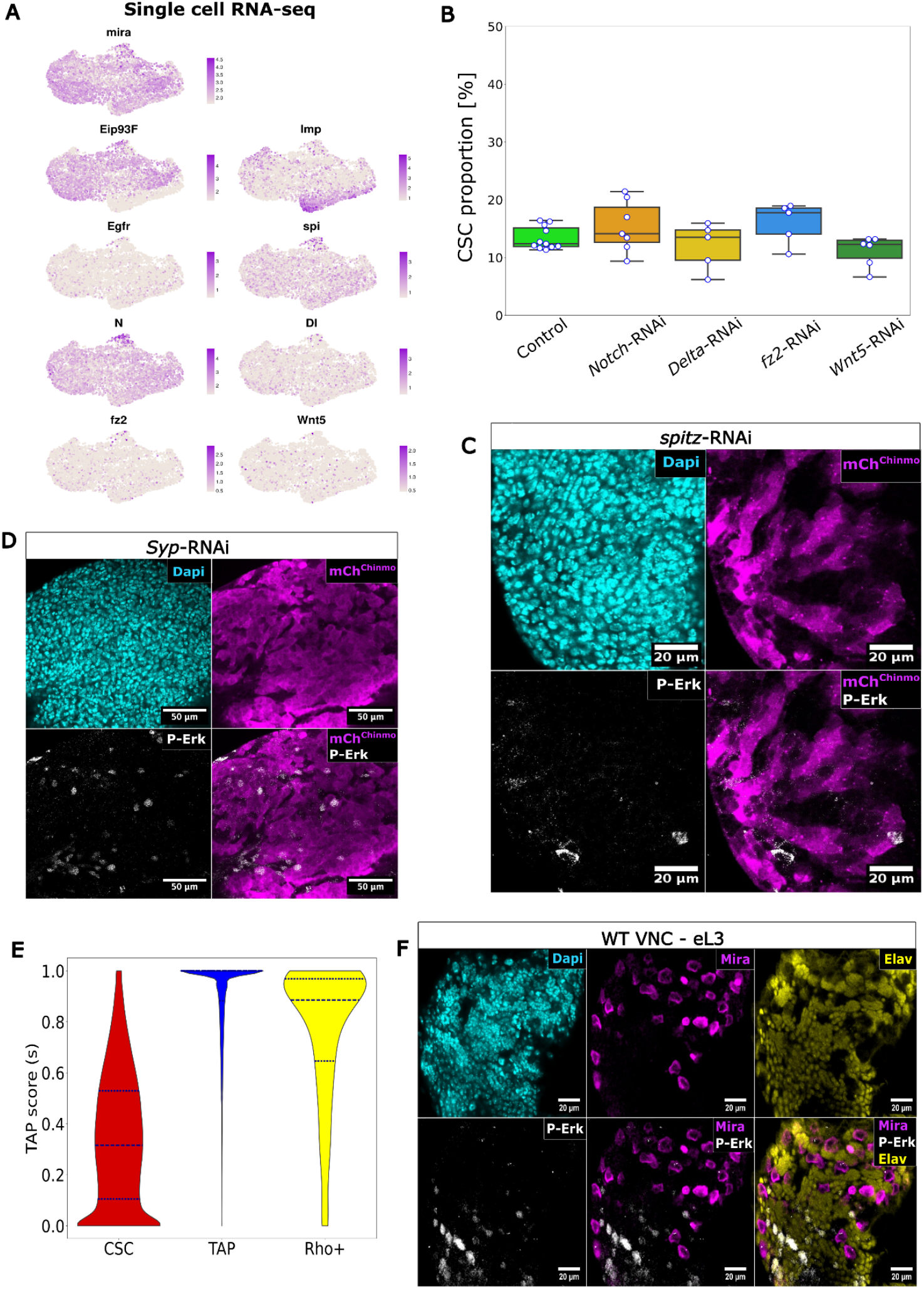
Molecular mechanisms involved in the CSC-to-TAP transition. (A) UMAPs of single cell RNA-seq data from *pox*^*n*^*>pros-RNAi* tumours in adults depicting expression of various genes including signaling pathways. (B) CSC proportions in tumours upon inhibition of components of the Notch and Wnt signaling pathways. (C,D) Dapi labels nuclei, mCh^chinmo^ labels CSCs. Absence of cytoplasmic P-Erk shows that EGFR pathway is not activated upon knockdown of *Spitz* or *Syp* in tumours. P-Erk antibody constitutively labels mitotic nuclei (see in E) (40x objective). (E) Rho+ cells exhibit a TAP score higher than CSCs but lower than TAPs indicating that they tend localize at the periphery of CSC clusters (around n = 150 000 cells in total, from 10 different tumours). (F) Around late L2/early L3, no P-Erk signal is observed in normal NSCs (Mira+) preceding the Imp-to-Syp transition, confocal section (40x objective)).

**Supplementary Figure S3:**
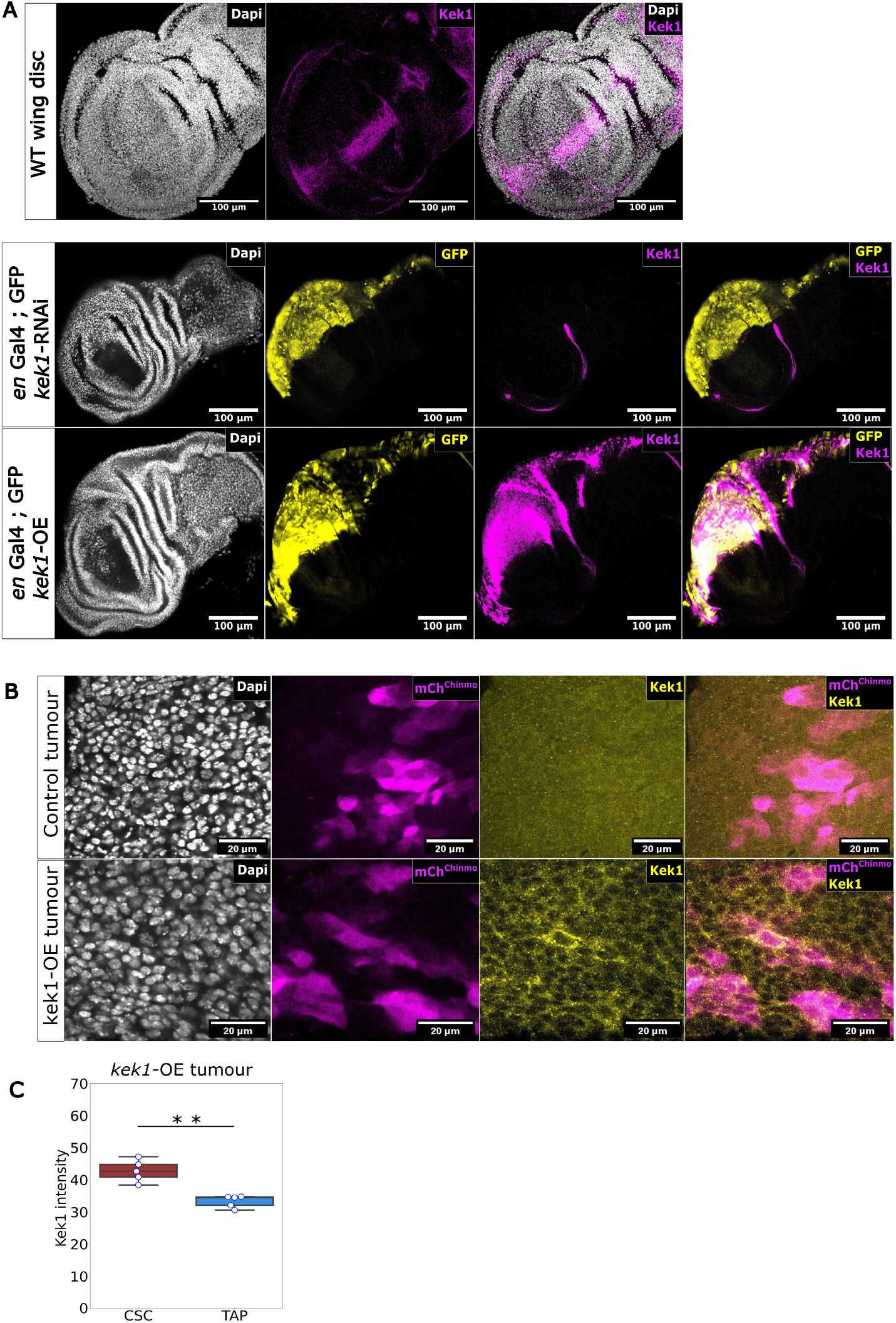
Characterization of the anti-Kek1 antibody and kek1-RNAi line. (A) Top panels -Anti-Kek1 reveals the wild type expression pattern of Kek1 in late L3 wing imaginal discs, as previously reported (Ghiglione et al., 2003). Middle panels - Kek1 conditional knockdown via RNAi in the posterior compartment using *en*-GAL4 leads to efficient silencing as detected by immunostaining. Bottom panels: Over-expression of *kek1* in the posterior compartment is strongly detected by the anti-Kek1 antibody. These experiments validate both the specificity of the newly generated anti-Kek1 antibody, and the efficiency of transgenic lines for knockdown and mis-expression of *kek1*. (B) Dapi labels all tumour nuclei. Top panels - The anti-Kek1 antibody does not detect Kek1 in chinmo+ CSCs. Bottom panels - The anti-Kek1 antibody detects Kek1 in tumour cells upon over-expression. (C) Quantification of Kek1 signal in CSCs vs TAPs upon *kek1* over-expression.

## Supplementary movies

**Supplementary Movie S1:** Timelapse 3D visualisation of a cell-intrinsic simulation (Figure 1F).

**Supplementary Movie S2:** z-stack of a cell-intrinsic 3D simulation outcome (Figure 1F).

**Supplementary Movie S3:** z-stack of a full tumour analysed with our segmentation and analysis pipeline. Spacing between slices is 4µm. Top right: DAPI. Top left: Imp (yellow) and Chinmo (Magenta). Bottom left: cell states. Bottom right: TAP score.

**Supplementary Movie S4:** z-stack of a cell-intrinsic 3D simulation outcome with *τ* = 1*d* and no differential adhesion.

**Supplementary Movie S5:** z-stack of a cell-extrinsic 3D simulation outcome with *τ* = 1*d* and differential adhesion (*ε*_*cc*_ = 9).

## Materials and Methods

### Key resources table

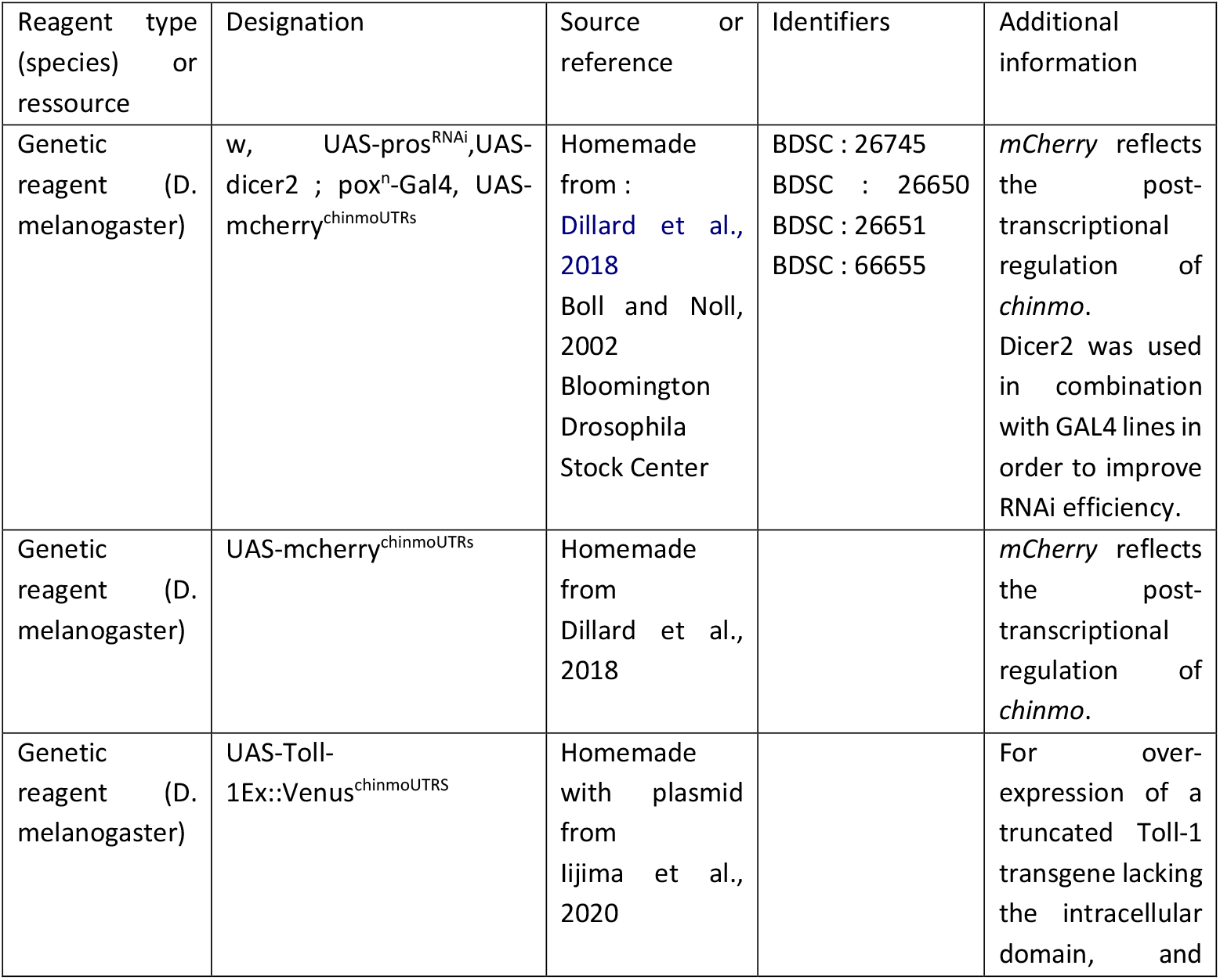
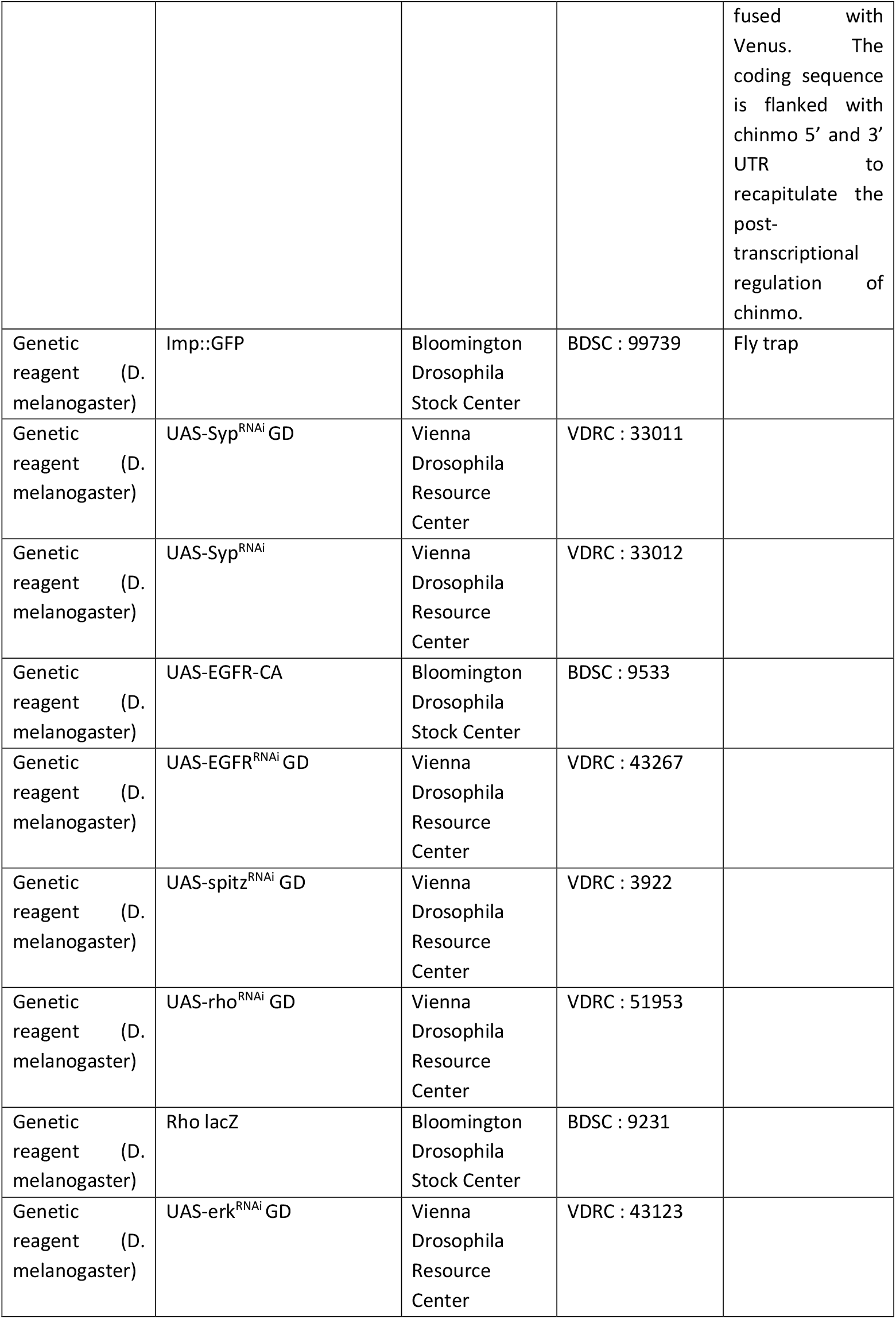
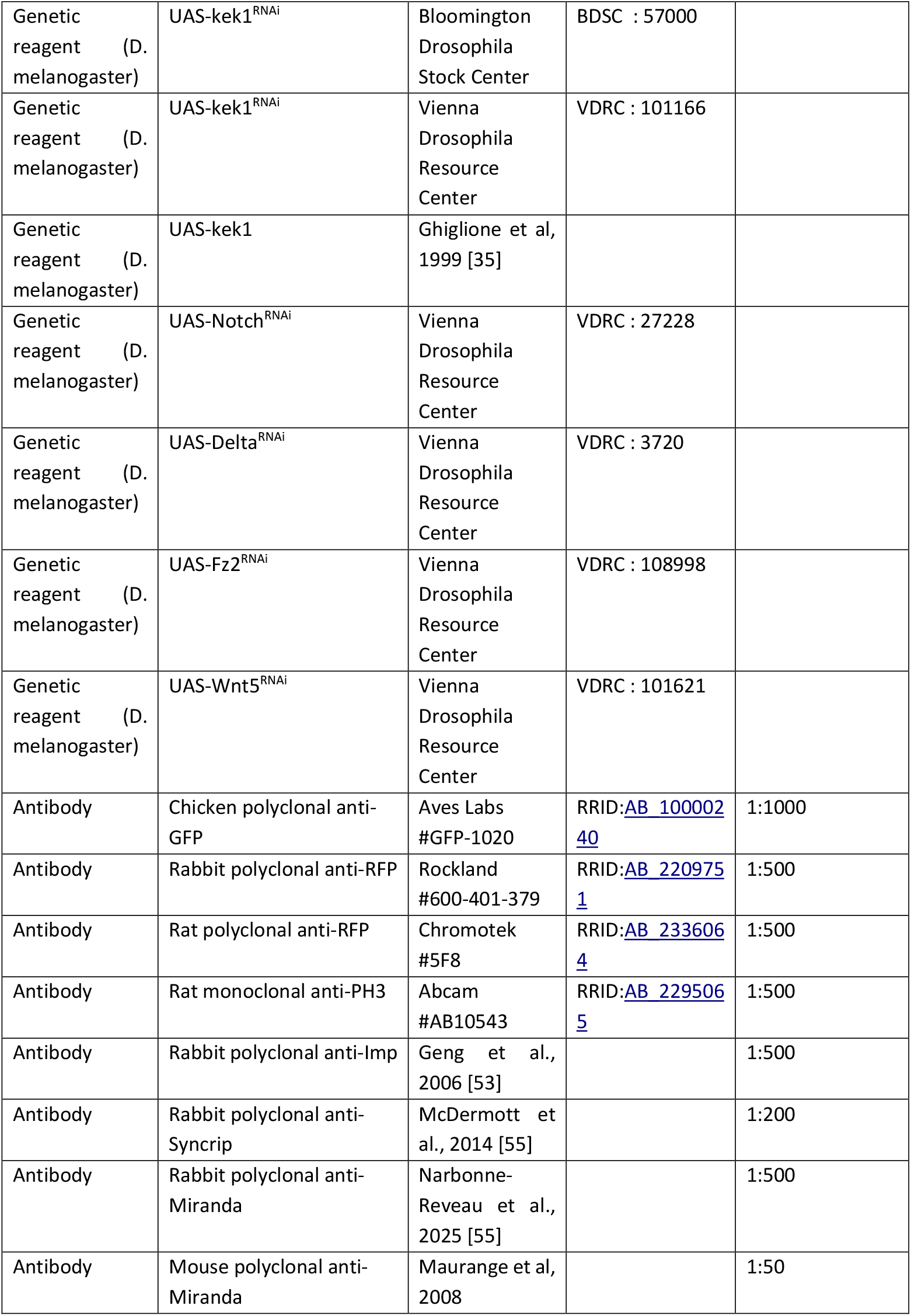
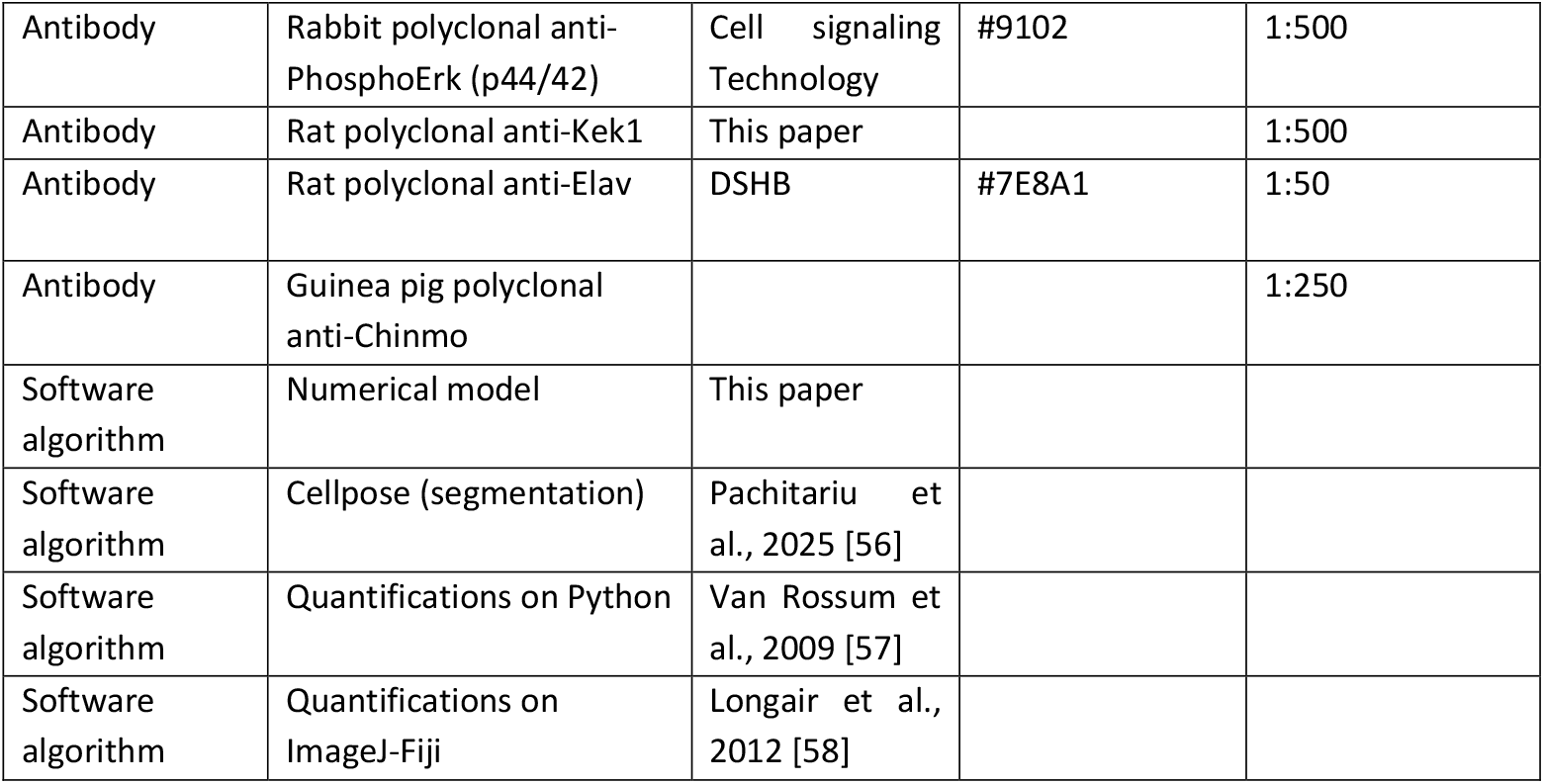

### Fly culture

Drosophila lines were kept at 18°C on standard medium (8 % cornmeal, 8 % yeast, 1 % agar).

### Fly lines

The *GAL4* line used are:

*pox*^*n*^*-GAL4* (active in six homologous thoracic neuroblasts of the ventral nerve cord (Boll and Noll, 2002).

The *UAS* lines used are:

-*UAS-mCherry*^*chinmoUTRs*^ (Dillard et al., 2018): The mCherry coding sequence is flanked by the 5’ and 3’ UTRs of chinmo mRNA reflecting the post-transcriptional regulation of endogenous chinmo mRNA. Consistently, we observe that mCherry always overlaps with endogenous Chinmo as detected by immunostaining (Gaultier et al., 2022; Genovese et al., 2019).

We used this approach to generate *UAS-Toll-1Ex::Venus*^*chinmoUTRS*^ transgenic line allowing specific expression of Toll-1Ex::Venus in CSCs.

Other stocks: *UAS-pros*^*RNAi1*^ (Bloomington Stock Centre #26745), *UAS-dicer2* (Bloomington Stock Centre #24650 and #24651) was used in combination with GAL4 lines in order to improve RNAi efficiency, *UAS-Toll1Ex::Venus*^*ChinmoUTRS*^ (Homemade with plasmid from Daiki Umetsu (Iijima et al., 2020), *UAS-EGFR-CA* (Bloomington Drosophila Stock Centre #9533), UAS-EGFR^RNAi^ (Vienna Drosophila RNAi Center #43267), *UAS-spitz*^*RNAi*^ (Vienna Drosophila RNAi Center #3922), *UAS-rho*^*RNAi*^ (Vienna Drosophila RNAi Center #51953), UAS-erk^RNAi^ (Vienna Drosophila RNAi Center #43123), *UAS-kek1*^*RNAi*^ (Bloomington Drosophila Stock Centre #57000), *UAS-kek1*^*RNAi*^ (Vienna Drosophila RNAi Center #101166), *UAS-kek1* (from Christian Ghiglione (Ghiglione et al., 1999)), *UAS-Notch*^*RNAi*^ (Vienna Drosophila RNAi Center #27228), *UAS-Delta*^*RNAi*^ (Vienna Drosophila RNAi Center #3720), *UAS-Fz2*^*RNAi*^ (Vienna Drosophila RNAi Center #108998), *UAS-Wnt5*^*RNAi*^ (Vienna Drosophila RNAi Center #101621).

The protein trap line used is *Imp-GFP* (Bloomington Stock Centre #99739). Other lines: *Rho-lacZ* (Bloomington Stock Centre #5467).

Fly crosses are set up and raised at 29°C.

### Generation of the UAS-Toll-1Ex::Venus^**chinmoUTRs**^ **plasmid and transgenic flies**

The Toll-1Ex::Venus coding sequence was cloned between the 5′ and 3′ untranslated transcribed regions (UTRs) of *chinmo* under the regulation of upstream activation sequences (UAS) using the In-Fusion® Snap Assembly Master Mix kit (Takara). The entry vector used was pUAS-mcherry^chinmoUTRs^ (Dillard et al., 2018) digested with PmeI to remove *mCherry* and to allow insertion of the Toll-1Ex::Venus sequence. The Toll-1Ex::Venus coding sequence was obtained from pUAS-Toll1Ex::Venus (a gift from Daiki Umetsu (Iijima et al., 2020)). The primers used were: pUAS-ChinReg_Toll-1ExV_iF GGTGCGGCCGCGTTTAAAC**ATGAGTCGACTAAAGGCCGCTTCC**; pUAS-ChinReg_Toll-1ExV_iR TCGGAAGTGGAGTTTAAAC**TTACTTGTACAGCTCGTCCATGCC**. Bold and underlines indicate sequences used for PCR amplification and overlapping sequences for In-Fusion, respectively.

Plasmid was sent to BestGene (https://www.thebestgene.com) for transgenesis via targeted insertion in attP-3B using Bloomington Stock #9748.

### Immunostaining

Larval and adult (5 days old) VNCs containing tumours were dissected in phosphate-buffered saline (PBS) and fixed for 30 min at room temperature (RT) in 4 % paraformaldéhyde/PBS. VNCs were rinsed with PBT (PBS containing 0.5 % Triton X-100) and incubated with primary antibodies for 5 days at 4°C. Secondary antibodies were incubated for 3 days at 4°C. VNCs were mounted in Vectashield (Clinisciences, France) without dapi for image acquisition. The following primary antibodies were used: chicken anti-GFP (1:1000, Aves #GFP-1020), rabbit anti-RFP (1:500, Rockland #600-401-379), rat anti-RFP (1:500, Chromotek #5F8), rat anti-PH3 (1:500, Abcam #AB10543), rabbit anti-Imp (1:500, Geng et al., 2006 [52]), rabbit anti-Syp (1:500, McDermott et al., 2014 [53]), rabbit anti-Miranda (1:500, Narbonne-Reveau et al., 2025 [54]), mouse anti-Miranda(1:500,), rabbit P-ERK (1:500, Cell signaling #9102), mouse anti-β-galactosidase (1:500, Promega #Z3781), rat anti-Elav (1:50, DSHB #7E8A10), guinea pig anti-Chinmo (1:250, gift from J. Enriquez).

### Production and validation of anti-Kek1 antibody

Anti-Kekkon1 polyclonal antibody was generated by Genscript (https://www.genscript.com/). The amino acid sequence used as epitope used for immunization is: MLVLTLLPGMILGTRYNQLHLYANGGASSSGPGGYRPAPSSQNEVYSIADSQPMTEDGYMPPSQHFPPTHSDLDPPAQQQSTCQTVCACKWKGGKQTVECIDRHLIQIPEHIDPNTQVLDMSGNKLQTLSNEQFIRANLLNLQKLYLRNCKIGEIERETFKGLTNLVELDLSHNLLVTVPSLALGHIPSLRELTLASNHIHKIESQAFGNTPSLHKLDLSHCDIQTISAQAFGGLQGLTLLRLNGNKLSELLPKTIETLSRLHGIELHDNPWLCDCRLRDTKLWLMKRNIPYPVAPVCSGGPERIIDRSFADLHVDEFACRPEMLPISHYVEAAMGENASITCRARAVPAANINWYWNGRLLANNSAFTAYQRIHMLEQVEGGFEKRSKLVLTNAQETDSSEFYCVAENRAGMAEANFTLHVSMRAAGMASLGSGQ.

The antibody was validated in wing discs compared to its known expression pattern, as well as with gain and loss of function conditions (Figure S2A).

### Image analysis

#### Images acquisition

Confocal image stacks were acquired using a Zeiss LSM880 microscope equipped with a 40× objective. Whole tumours were imaged in 3D using z-stacks with 1 μm intervals between optical sections.

#### Volumes quantification

tumour volumes and CSC proportions were quantified using FIJI (ImageJ). Binary masks of tumour tissue and CSC clusters were generated through intensity thresholding. Volumes were subsequently measured using the MorphoLibJ plugin based on voxel counting.

#### Tumour segmentation

Cell segmentation was performed using Cellpose-SAM (Pachitariu et al., 2025), employing the pre-trained models for nuclear (DAPI) and cytoplasmic (Imp and P-Erk) labeling. This allowed the segmentation of individual cells within the densely packed tumour tissue and the classification of cells based on marker expression. To prevent erroneous merging of adjacent cells during downstream quantification, a 2-pixel erosion was applied to all segmented masks.

#### ROIs extraction

Quantifications post segmentation were made on FIJI (ImageJ) to manually correct the segmentation (Schindelin et al., 2012). Regions of interest (ROIs) corresponding to individual cells were extracted using the “Analyze Particles” function, providing information such as the XYZ coordinates and mean fluorescence intensity for each marker.

#### Data processing and classification

Data processing and cell classification were performed using Python.

#### Segregation Index (*μ*) (Figure 1H)

In all quantifications, CSCs are identified using the *mCherry::chinmoUTRs* intensity. Chinmo+ cells are classified as cancer stem cells (CSC). Chinmo-cells are classified as Transit Amplifying Progenitors (TAPs, thus also including NPCs). For each cell, the segregation index (μ) is calculated based on the identity of the neighbouring cells. We use a distance-based criterion for neighbourhood: cells in a 7µm sphere around the target cell are considered as neighbours. μ is normalized between -1 for perfect mixing and +1 for perfect segregation. μ=0 corresponds to a balance of CSC and TAPs that matches the tumour-level proportions. Thus, for a cell with the *x* identity (either CSC or TAP), we have:

If *q*_*x*_ ≥ *Q*_*x*_

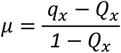

If *q*_*x*_ < *Q*_*x*_

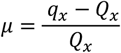

Where *q*_*x*_ is the proportion of neighbours with the *x* identity, and *Q*_*x*_ is the tumour-level proportion of cells with the *x* identity.

#### TAP score (s) (Figures 2D)

In all quantifications, CSCs are identified using the mCherry and Imp intensities obtained after immunostaining. Chinmo^+^ Imp^+^ cells were classified as CSCs, Chinmo^−^ Imp^+^ cells as tTAPs, and Chinmo^−^ Imp^−^ cells as TAPs. For each cell, the TAP score *s* is calculated based on the identity of its neighbouring cells in 3D. Again, cells in a 7µm sphere around the target cell are considered as neighbours.

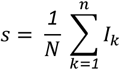

For each cell, *N* is the number of neighbours, and the contribution *I*_*k*_ of neighbour *k* depends on its identity. For CSC neighbours, *I*_*k*_ = *0*, for TAPs, *I*_*k*_ = *1* and for tTAPs, *I*_*k*_ = *0*.*5*. Consequently, a cell only surrounded by CSCs has a score of 0, whereas a cell only surrounded by TAPs has a score of 1.

### Statistical analysis

For each volume or transition rule experiments, at least 4 tumours were quantified. To test the significance of volume or proportion differences between conditions, we performed a Mann-Whitney test. To test the significance of TAP distribution differences, we performed a Student test. The statistical analysis was performed using the Scipy Python library. Results are represented as boxplots or violin plots using the seaborn or matplotlib Python libraries. Sample sizes are reported in the figures.

### Numerical model

#### Rationale of the model

We developed an agent-based Monte-Carlo stochastic numerical model. This approach enables rapid and systematic testing of different differentiation scenarios used to motivate experiments. Conversely, experimental findings can be recapitulated and interpreted within the model framework.

Our model simulates 3D tumour development driven by two distinct cell types: Chinmo^+^ cells (C), representing cancer stem cells (CSCs), and Syncrip^+^ cells (S), corresponding to transit-amplifying progenitors (TAPs). Starting from an initial configuration at time t = 0 with a tuneable number of cells, the simulation progresses up to a final time fixed by the user. Throughout the simulation, cells can undergo division, change identity (C→S), enter a non-proliferative state, and move to account for steric repulsion when new cells are generated. The rates governing stochastic division and transition events can be modulated by the user and depend on the scenario tested.

Cell–cell physical interactions are governed by a Lennard-Jones potential to account for steric constraints and dense cell packing. The Lennard-Jones potential sets an effective, preferred distance between cells, typically corresponding to their own size. The interaction strength, tuned by the potential depth, can either be the same for all cells (*ε*_*cc*_ = *ε*_*cs*_ = *ε*_*ss*_) or heterogeneous, enabling modelling of differential affinity/adhesion between cell types.

#### Model Parameters and consistency with (Genovese et al., 2019)

The model presented here requires setting a few key parameters. Importantly:

- The division rates *k*_*c*_ and *k*_*s*_ of C and S cells.
- The rate of cell-cycle exit *k*_*cce*_ of S cells.
- The C→S transition rate *k*_*t*_, which can be the same for all CSCs or depend on their neighbourhood, depending on the scenario tested (intrinsic or extrinsic).
- The interaction potential strengths *ε*_*cc*_, *ε*_*cs*_ and *ε*_*ss*_ between cells. Unless we specifically want to test the role of differential affinity/adhesion, we set *ε*_*cc*_ = *ε*_*cs*_ = *ε*_*ss*_ = 1.

Note that *k*_*c*_, *k*_*s*_, *k*_*cce*_ and the average transition rate < *k*_*t*_ > (used for the intrinsic scenario, see below) are directly extracted from our previous study using clonal analysis (Genovese et al., 2019), in which we quantified transition and cell-cycle exit probabilities per division cycle. Here we will detail the consistency between the rates used here and the parameters inferred in (Genovese et al., 2019).

In the clone model, we obtained the division times *T*_*c*_ = 1/*k*_*c*_ = 1.6*d* and *T*_*s*_ = 1/*k*_*s*_ = 1.3*d*. We also inferred the average probabilities for C cells to undergo differentiation during a division cycle. The probability for a C cell to self-renew upon division was found to be *p*(*C* →*CC*)=0.64, the probability to give rise to two S cells *p*(*C*→*SS*)=0.21 and the probability to give rise to one cell of each type *p*(*C*→*CS*)=0.15. In this paper, we chose to reformulate these probabilities in terms of a single average transition rate < *k*_*t*_ > per unit time. This was done for multiple reasons. First, it makes the model more parsimonious, second, we have no a priori information on a link between transition and division, and third, experimentally we only have access to the probability *p*_*t*_ (*s*) for single cells to be undergoing transition, which directly related to the transition rate. Equating the expected number of new S cells per C cell division in both formulations, we obtain that:

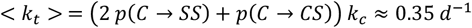

We use a similar approach to obtain the rate of cell cycle exit of S cells, *k*_*cce*_. Equating the rate of creation of non-proliferative cells in both formulations, we find that *k*_*cce*_ *S* = *q*_*s*_ (2*k*_*s*_ *S* + *k*_*t*_*C*), such that:

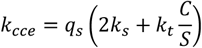

Where *q*_*s*_ = 0.47 is the probability for a new S cell to be non-proliferative in the clone model, C is the number of C cells, and S the number of S cells. When the system reaches stable proportions, C*/S* is constant and C and S populations have the same growth rate, such that *d*_*t*_*C* = (*k*_*c*_ − *k*_*t*_)*C* = *λC* and *d*_*t*_*S* = (*k*_*s*_ − *k*_*cce*_)*S* + *k*_*t*_*C* = *λS*, which leads to:

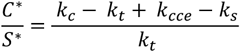

Combining the above equations, we find:

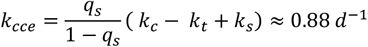

#### Simulation workflow

Once these parameters are defined, the simulation proceeds through the following steps, repeated at each time step (we set the time step *T* to 1 hour in the simulations):

#### 1. Cell division and cell cycle exit

C and S cells undergo probabilistic division, and S cells undergo cell-cycle exit, based on their respective rates (see above). For each cell, the probability that division or cell cycle exit occur during one time step of duration *T* is:

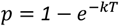

Where k corresponds to the rate of the considered event (*k*_*c*_, *k*_*s*_, *k*_*cce*_). When a division event occurs, the two resulting daughter cells positions are initialized at a distance *r*_*0*_/10, where *r*_*0*_ is the cell radius, centered around the position of the mother cell and at a random angle. This is followed by the energy minimization step.

#### 2. Movement through energy minimization

New cells generated in the bulk push on the surrounding cells. In the simulations, the steric interactions between cells are mimicked by the Lennard-Jones potential, and we use global energy minimization to calculate cells’ displacement at each time step. The energy *E* is calculated by summing the Lennard Jones potentials *V* of all possible N interactions between cell pairs, such that:

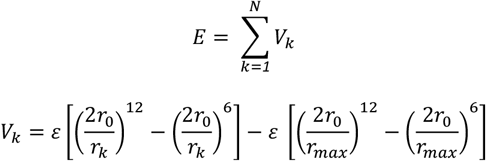

With *ε* the interaction strength, *r*_*0*_ the prescribed cell radius, *r*_*k*._ the distance between the 2 cells of cell pair *k* and *r*_*max*_ the maximum distance considered (since the potential vanishes at long distance, we only consider cells within a certain range to compute the total energy).

Energy minimization is performed using a Monte Carlo–type relaxation scheme. Each sweep consists of *N*_*cells*_ single-cell displacement attempts, where at each attempt a cell is selected at random, ensuring statistically uniform sampling of all cells. The proposed displacement combines a deterministic force-driven term (local energy gradient) and a Gaussian random term. A displacement is always accepted if it lowers the energy, and accepted with probability *e*^−Δ*E*^ if it increases it (Metropolis criterion). The process is iterated over successive sweeps until convergence is reached. Convergence is reached when *N*_*cells*_10 consecutive displacement attempts produce sufficiently small energy variations, ensuring that the criterion scales with system size. Variations are considered small enough if *ΔE*^2^/*N*_*cells*_< 10^−4^, i.e. the energy change per cell is below a threshold independent of system size.

#### 3. Cell rearrangements via swapping for differential adhesion

Some of our simulations were designed to specifically test the effect of increased segregation, without changing the transition rules (Figure 3E,F). As discussed earlier, this is done by increasing the interaction strength *ε*_*cc*_ among CSCs. A common strategy to implement differential affinity or adhesion in agent-based models is to allow neighbouring cells with different identities to swap positions if it lowers the energy minimum. Note that in real cells, sorting also requires motility. Thus, we implemented test-swapping at a rate *k*_*swap*_. For each cell, the probability that it is chosen for test-swapping during one time step of duration *T* is thus given by:

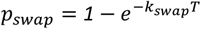

To determine with which neighbour swapping occurs, the post-swap energy is computed for all possible swaps. The neighbour that lowers the overall energy the most is chosen for swapping. If none of the tested swaps lower the energy, no swapping occurs. Of note, this ensures that no swapping happens if *ε*_*cc*_ = *ε*_*cs*_ = *ε*_*ss*_.

#### 4. C→S transition

Cell-intrinsic scenario (Figure 1F,G): In the cell-intrinsic transition scenario, all CSCs have the same transition rate *k*_*t*_ such that for all CSCs:

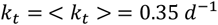

and the probability for a CSC to initiate transition at each simulation time step of duration T is:

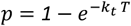

Cell-extrinsic scenario (Figure 3A-F): Our TAP score analysis revealed that the probability to transit is a quantifiable function of neighbourhood identity. Hence, we devised simulations where the probability to transit is determined locally for each cell. Specifically, we measured experimentally the proportion of CSCs *currently* undergoing transition, *p*_*t*_(*s*). This observed proportion depends on the transition rate *k*_*t*_ (*s*) but also on the time *τ* spent by CSCs in the transition state, such that the transition rate we input in simulations is *k*_*t*_ (*s*) = *p*_*t*_ (*s*)/*τ* (Little’s law of occupancy). For *p*_*t*_ (*s*), we directly use a sigmoid fit of the experimental curve. We conducted sensitivity analyses for *τ*, and show the simulations outcome for the resulting *k*_*t*_ (*s*). The probability for a transition to occur in a given CSC with TAP score *s* at each simulation time step is:

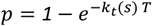

## Data availability

Codes generated for this study are available at: codes will be made available for publication Single-cell data used in this study were already published (Genovese et al., 2019) and are available at **NCBI Gene Expression Omnibus** ID GSE114986

## Author contributions

EL performed the experiments, image analysis, quantifications, and simulations. CR designed the model with RC, and wrote the 3D simulation code. LB created the Toll-1 constructs. RC et CM conceived and supervised the project. EL, RC and CM discussed the data, quantifications, and model, and wrote the paper.

## Acknowledgements

We warmly thank many colleagues from the Drosophila community for flies, plasmids and antibodies. We are grateful to Julien Leclercq for help on single-cell analysis. Funding to CM was supported by La Ligue Contre le Cancer (Equipe Labellisée). EL was supported by Centuri and Fondation pour la Recherche Médicale (FRM), and CR was supported by Centuri. This work was also supported by a government grant managed by the Agence Nationale de la Recherche under the France 2030 program, with the reference number ANR-24-EXCI-0001, ANR-24-EXCI-0002, ANR-24-EXCI-0003, ANR-24-EXCI-0004, ANR-24-EXCI-0005. We also received help from the Centre South-ROCK (INCa-Cancer_18695). Stocks obtained from the Bloomington Drosophila Stock Center (NIH P40OD018537), Vienna Drosophila Resource Center (VDRC) and the Drosophila Genomics Resource Center were used in this study. We also thank Developmental Studies Hybridoma Bank (DSHB). We are grateful to the imaging facility at IBDM, member of the National Infrastructure France-BioImaging (https://ror.org/01y7vt929) supported by the French National Research Agency (ANR-24-INSB-0005 FBI BIOGEN).

