## Supplementary figures and images for "System-level regulation of hierarchical transitions in a tumour lineage"

### Supplementary Movie S1

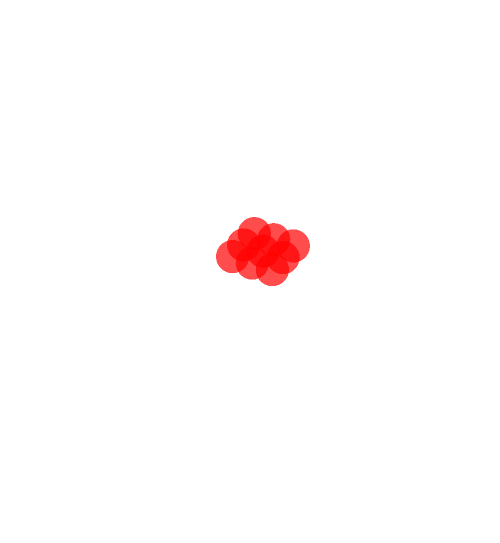

### Supplementary Movie S2

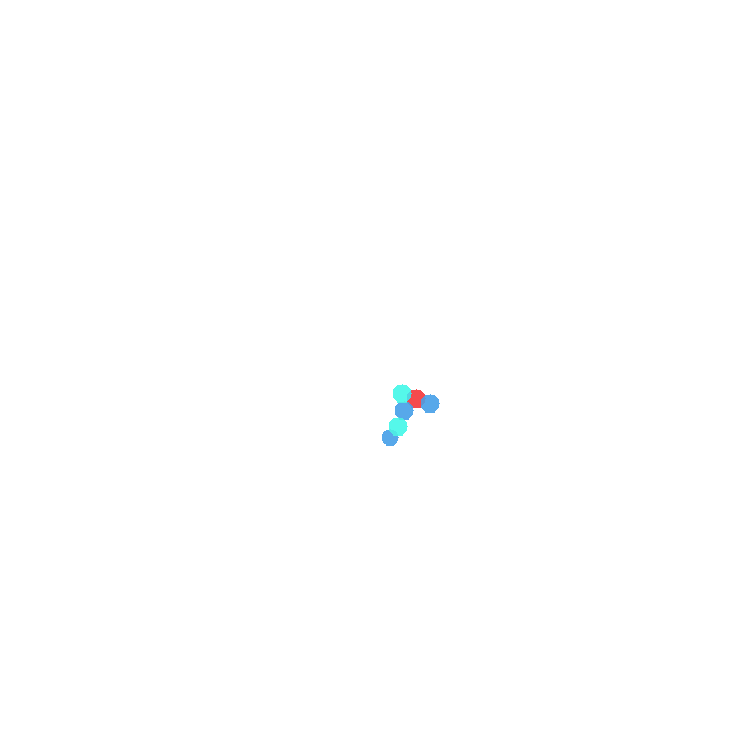

### Supplementary Movie S3

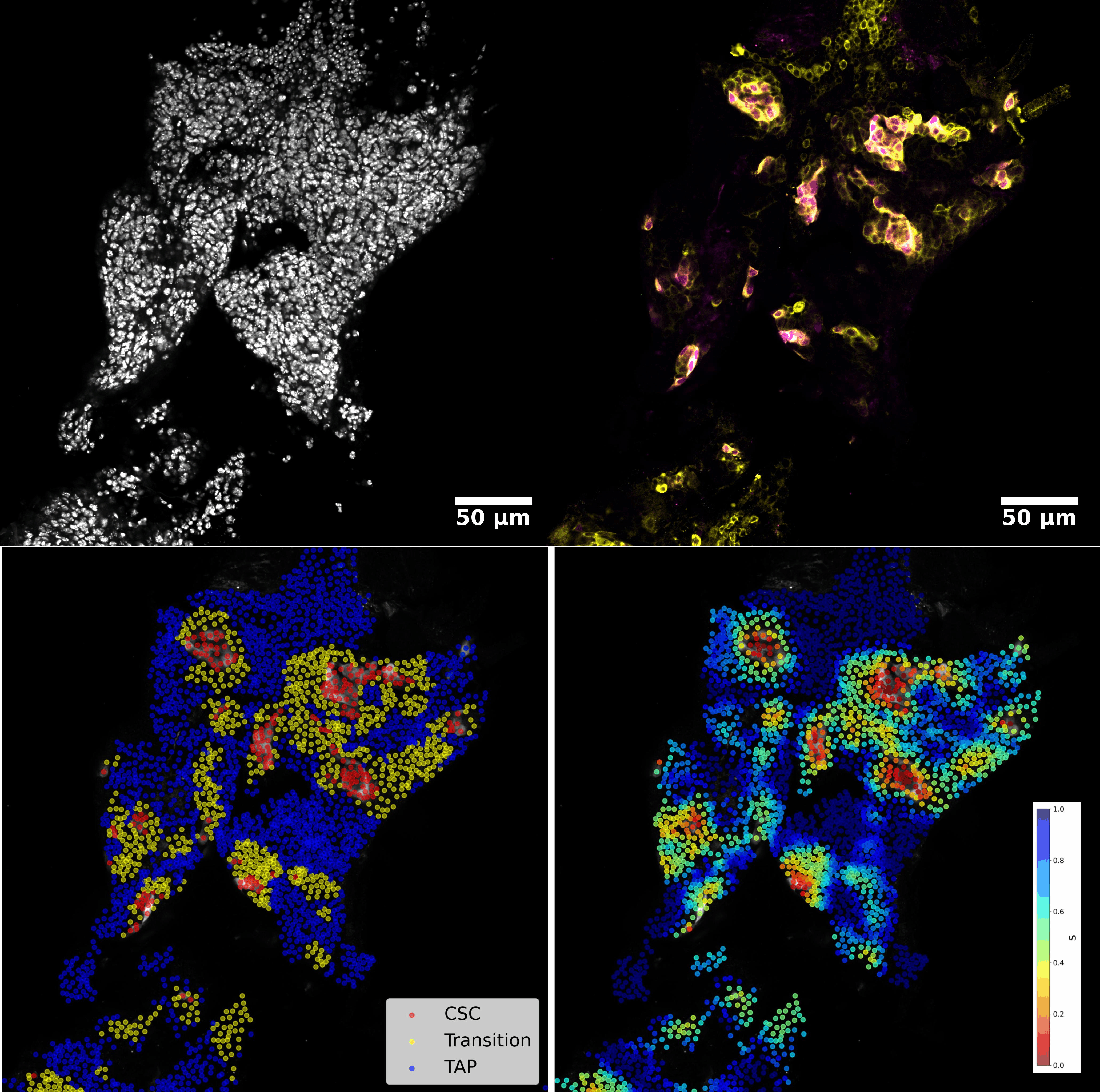

### Supplementary Movie S4

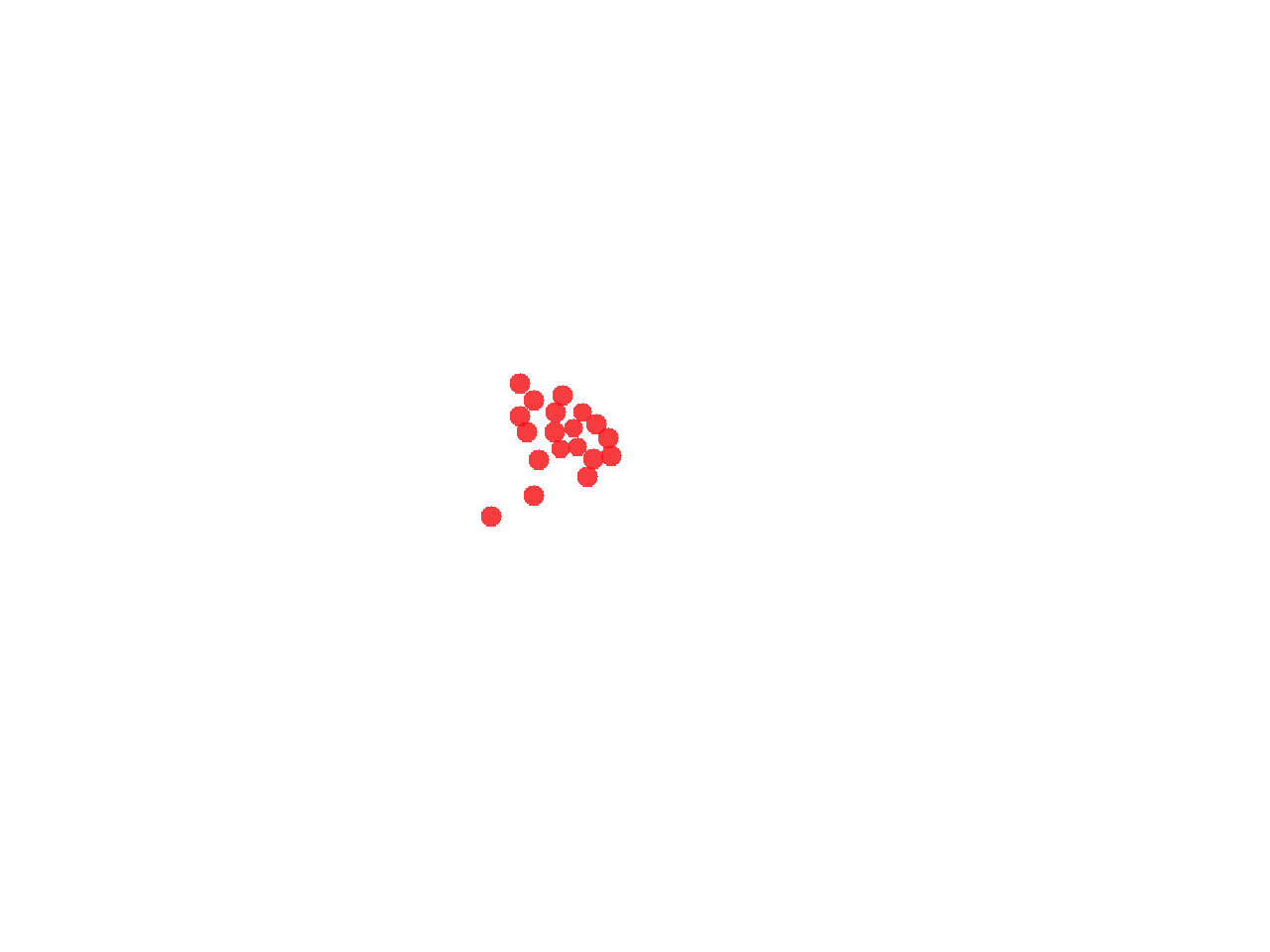

### Supplementary Movie S5

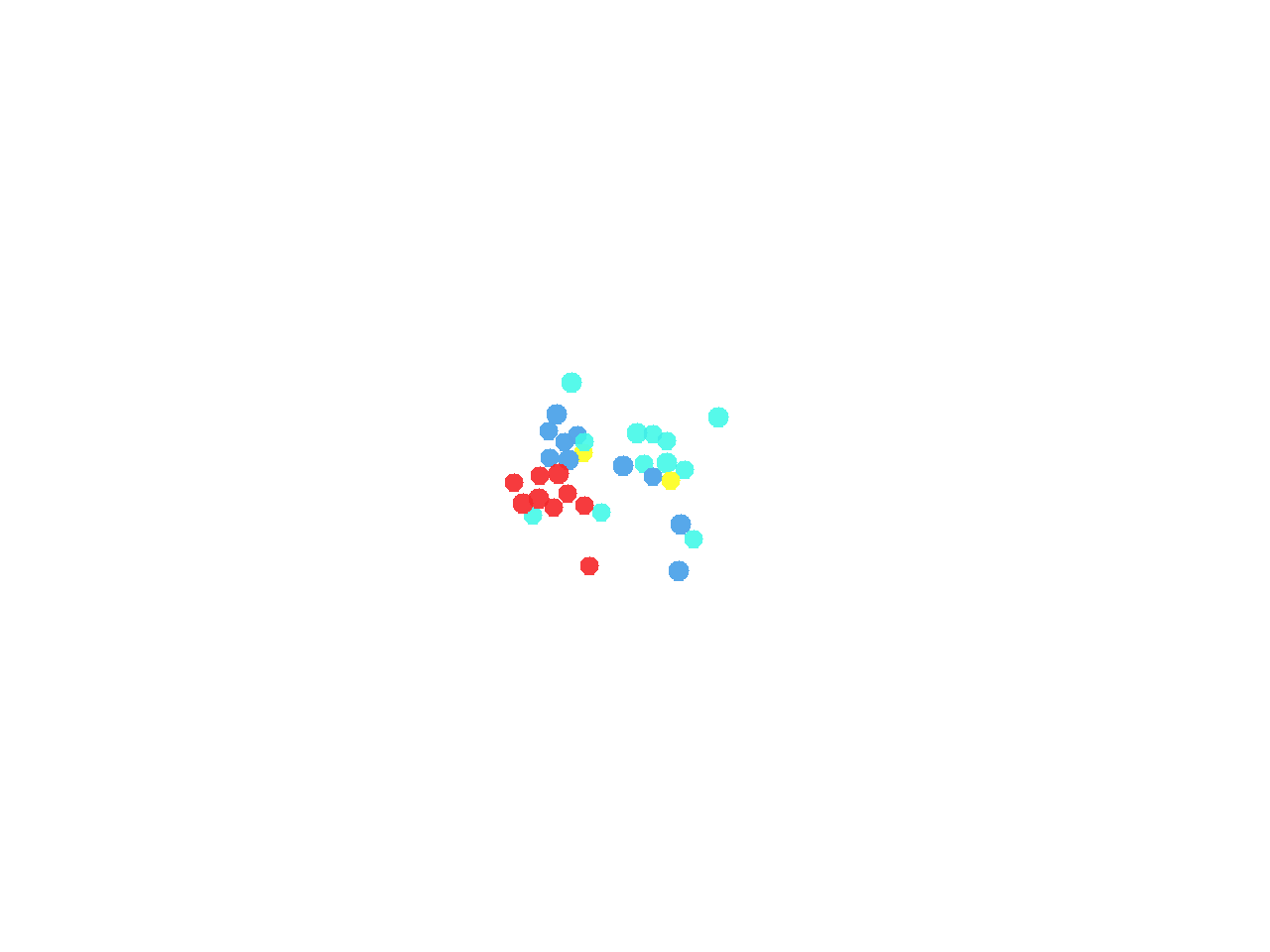
